# INTEGUMENT-SPECIFIC TRANSCRIPTIONAL REGULATION IN THE MID-STAGE OF FLAX SEED DEVELOPMENT INFLUENCES THE RELEASE OF MUCILAGE AND THE SEED OIL CONTENT

**DOI:** 10.1101/2021.09.09.459608

**Authors:** Fabien Miart, Jean-Xavier Fontaine, Gaëlle Mongelard, Christopher Wattier, Michelle Lequart-Pillon, Sophie Bouton, Roland Molinié, Nelly Dubrulle, Françoise Fournet, Hervé Demailly, Romain Roulard, Loïc Dupont, Arezki Boudaoud, Brigitte Thomasset, Laurent Gutierrez, Olivier Van Wuytswinkel, François Mesnard, Karine Pageau

**Affiliations:** Laboratoire de Biologie des Plantes et Innovation, UMRT 1158 INRA BioEcoAgro, UPJV, UFR des Sciences, 33 rue St Leu, F-80039, Amiens, France; Centre de Ressources Régionales en Biologie Moléculaire, UPJV, UFR des Sciences, Bâtiment Serres-Transfert Rue Dallery - Passage du sourire d’Avril, F-80039, Amiens, France; Laboratoire Joliot Curie, CNRS, ENS Lyon, UCB Lyon 1, Université de Lyon, 46 Allée d’Italie, F-69364, Lyon Cedex 07, France; Laboratoire de Reproduction et Développement des Plantes, INRA, CNRS, ENS Lyon, Université de Lyon, 46 Allée d’Italie, F-69364, Lyon Cedex 07, France; Laboratoire de Réactivité et de Chimie des Solides, UMR CNRS 7314, UPJV, UFR des Sciences, 33 rue Saint Leu, F-80039, Amiens, France; Laboratoire de Génie Enzymatique et Cellulaire, UMR CNRS 7025, UTC, Université de Technologie de Compiègne, F-60205, Compiègne, France

**Keywords:** Carbon partitioning, flax inbreed lines, GA metabolism, mucilage, fatty acids, 25 DAF

## Abstract

Flax (*Linum usitatissimum* L.) seed oil, which accumulates in the embryo, and mucilage, which is synthesized in the seed coat, are of great economic importance for food, pharmaceutical as well as chemical industries. Theories on the link between oil and mucilage production in seeds consist in the spatio-temporal competition of both compounds for photosynthates during the very early stages of seed development. In this study, we demonstrate a positive relationship between seed oil production and seed coat mucilage extrusion in the agronomic model, flax. Three recombinant inbred lines were selected for low, medium and high mucilage and seed oil contents. Metabolite and transcript profiling (1H NMR and DNA oligo-microarrays) was performed on the seeds during seed development. These analyses showed main changes in the seed coat transcriptome during the mid-phase of seed development (25 Days Post-Anthesis), once the mucilage biosynthesis and modification processes are thought to be finished. These transcriptome changes comprised genes that are putatively involved in mucilage chemical modification and oil synthesis, as well as gibberellic acid (GA) metabolism. The results of these integrative biology approach, suggest that transcriptional regulations of seed oil and fatty acid (FA) metabolism could occur in the seed coat during the mid-stage of seed development, once the seed coat carbon supplies have been used for mucilage biosynthesis and mechanochemical properties of the mucilage secretory cells.

## INTRODUCTION

Seeds development involves complex regulatory networks from the genomic to the phenomic scales which affect seed growth, quality and germination process [1,2,3,4]. Seeds consist of three major tissues, i.e. the embryo, the endosperm and the seed coat, each of them differing from their genetic origin and chemical composition. Oilseed species accumulate oils in their embryo as a major storage compound, and mucilage in the most external epidermal cell layer [4–6]. For flax seed, lignans are also localized in the outer integument [7,8]. Lignans, seed oil and mucilage are of great importance in the emerging bioeconomy [9], for fundamental and applied research on the cell wall polysaccharides structure and function [10,11]], for human health [12–15] as well as for improving green chemistry towards various industrial applications including cosmetic and food products (reviewed in [16]). Mucilage of flax contains RG I (rhamnogalacturonans type I) with an important fraction of highly branched AXs (Arabinoxylans, [17–19]). These two main polysaccharidic fractions are deposited in the Mucilage Secretory Cells (MSCs) as successive layers. RG I is positioned at the upper face of MSCs while AXs and small amount of glucans are found at their bottom side. A mixture of both layers is positioned at their interface, constituting a supplementary composite layer [20]. Moreover, the flax MSCs internal structure lacks the central volcano-shaped cellulosic structure called the columella which is typical of Arabidopsis MSCs [7,21]. Flax seed is also the main source of lignans in the form of complexed macromolecule in plants, among which secoisolariciresinoldiglucoside (SDG) is the most abundant [22]. In flax seed, SDG is incorporated into the lignan macromolecule via a hydroxy-methylglutaric acid (HMG) ester linkage [23]. The composition of polyphenols in the lignan macromolecule may vary and coumaric acid glucoside (CAG), ferulic acid glucoside (FAG) and herbacetin diglucoside (HDG) can be linked on SDG residue [23,24]. Flax seeds also have a high oil content with 96% of TAGs (triacylglycerols) mainly localized in the embryo. The major fatty acid found in the flax seed oil is linolenic acid (ω-3, LIN, C18:3, 50-62%) and palmitic acid (PAL, C16:0, 4-6%), steric acid (STE, C18:0, 2-4,4%), oleic acid (OLE, C18:1, 10-24,2%), linoleic acid (ω-6, LIO, C18:2, 12-18%) are also present [25–27]. Oilseed species, such as Arabidopsis and flax, accumulate oil from early-middle to late stages of seed development [5,28].

Several ways have been proposed to improve seed storage through modifications of the carbon-nitrogen flows [29–32], for example modifications of key steps in the mucilage biosynthesis pathway in the seed coat to support accumulation of seed oil in the embryo [33]. Moreover, recent studies on the model plant Arabidopsis and on *Medicago orbicularis*, have demonstrated negative correlations between seed coat compounds accumulation, including mucilage and proanthocyanidins, and the oil content in the embryo [31,33–35]. Interestingly, some proteins and transcription factors like the multi-media complex WD40-bHLH-MYB (TTG1-EGL3-MYB5) are involved in the control of mucilage and phenylpropanoids synthesis [31,36–40]. As an example, it has been shown MYB5 transcription factor acts synergistically with the TT2 factor on the control of mucilage biosynthesis pathway but also plays a minor role in the synthesis of proanthocyans (Pas) [38,40,41]. Finally, the analysis of different transparent testa mutants evidenced that a significant decrease in PAs in the seed coat is correlated with a significant increase in fatty acid content in embryo [31,42].

In flax, despite the reported of negative connections between seed coat-associated phenylpropanoids (SDG and hydroxycinnamic acid glucosides) and ω-3 content revealed by 1^H^-NMR-metabolomics analyses [43], to our best knowledge, so far no correlations have been demonstrated between mucilage and oil or fatty acid (FA) content [44,45]. Lack of such correlations could be explained by the complex chemical composition of the flax mucilage but also because of various extraction methods that exists to characterize mucilage [17,46]. Current hypothesis on seed coat/endosperm (SCE) and embryo metabolic interactions consists in the temporal and differential allocation of the main carbon sources, i.e. sucrose, towards mucilage synthesis and redirection of the unused organic molecules towards oil production in the embryo through the apoplastic route which is restricted to the globular embryo stage [33,47]. An illustration of this was given by the observation of an increased embryo oil content in the Arabidopsis *glabra2* mutant. *GLABRA2* codes for a transcription factor which controls positively the seed expression of *MUM4* gene coding for a rhamnose synthase required for mucilage biosynthesis. Thus, it was proposed that increased seed oil content in the *glabra2* mutant was caused by the re-allocation of carbon to the embryo because of the decreased synthesis of mucilage [33].

Perception of mechanical pressure has also been proposed as a possible way to explain regulations of the SCE-embryo interactions. Recently, it was shown that the regulation of the seed size was shown to be more complex than expected and depends on a fine modulation between the expansion of the albumen and that of the seed coat, all controlled by the physical properties of the cell wall of these two tissues and by a dependent Ca^2+^ signaling cascade [48– 50]. The expression of the *ELA1* seed size regulator, which contributes to the control of the gibberellic acid (GA) metabolism and consequently affects the seed size, was found to be regulated by mechanical forces from embryo and endosperm pressures exerted on the seed coat [50]. GA metabolism is connected to the cell wall properties, mucilage, starch, FAs metabolism and seed growth [42,50–54].

Engineering seed coat mucilage and oil production using biotechnological approaches, therefore, requires a better understanding of the spatio-temporal metabolic and transcriptomic regulations of both biosynthetic pathways. This work specifically focused on the regulation of the mucilage release, GA metabolism as well as analysis of the main carbon source unloading and partitioning. Flax was chosen as a model crop species due to its similarity to the fundamental model *Arabidopsis thaliana* according to mucilage and seed oil importance and as a growing oilseed crop for its high bioeconomic importance. Using two flax recombinant inbred lines (RILs) selected for opposite phenotypes in the mucilage release and oil content compared with a reference line, i.e. RIL 44 [20], we show that the main phenotypic differences originate more likely from the transcriptomic level rather than the metabolic level and occur especially in the seed coat in the mid-phase of flax seed development (25 Days Post-Anthesis, DPA). Similar spatio-temporal transcriptional regulation has been observed for GA metabolism, which fits well with differences in the mechanical forces exerted from the embryo on the SCE, physical properties analysis of the seed coat surfaces and the seed size and shape. Our results suggested that from the early to the mid-stage of flax seed development, the main sugar supplies are especially used in the seed coat for the mucilage biosynthesis, modifications and cell wall reinforcement, affecting the mechanochemical properties of the MSCs and their sensitivity to water imbibition [20]. Hereby, we propose that during the mid-stage of seed development, oil, mucilage and GA metabolism-related genes would be regulated by the coordinated use of carbon and by the mechanical forces induced by the expansion of the embryo on the seed coat and endosperm. All these metabolic pathways would modulate in turn the FA production in the embryo and the seed growth.

## MATERIALS AND METHODS

### Plant Materials and Growth Conditions

Breeder seeds of *Linum usitatissimum* L. cv Oliver and Viking and the recombinant inbred lines (RIL) population of 186 individuals were provided by GIE Linea Semences de Lin, Grandvilliers, France. Seeds were selfed in greenhouse up to F8 and multiplied in the field up to F12. Plants of Oliver, Viking, RIL 44, RIL 283 and RIL 80 from generation F12 were grown in soil in a greenhouse under the following conditions: 60 % humidity, 21 °C/15 °C day/night regime and with a 16-h photoperiod. For biological replicates, both parental cultivars and the selected RILs were harvested in three different greenhouses under the same growth conditions. The position of the plants in each greenhouse was interchanged in order to reduce light intensity or greenhouse positional effects. For analysis of the RILs, heads were tagged at anthesis. Developing seeds were collected at different stages of development and then dissected. The embryos and seed coat/endosperm (SCE) of each seed were isolated and then frozen in liquid nitrogen.

### Plant Phenotyping

The RILs population grown in the field was phenotyped in parallel on the F10 generation for mucilage released content and seed oil content and FA composition. Measurements of the mucilage released content were performed by carefully placing seeds along a line on 6 % (*w*/*v*) agarose plates containing 0.004 % (*w*/*v*) of the dye toluidine blue O, with five seeds per RIL and six RILs per plate. Two biological replicates were analyzed for each RILs on two different plates (20 seeds per RILs). After 24 h of mucilage release, the plates were placed on a light box and a picture was acquired using a standard high-resolution digital camera and a fixed camera mount.

For analysis of the released-mucilage content of the three selected RILs, the area of released-mucilage and area of the corresponding seed were manually segmented and quantified using the “colour threshold” function in the ImageJ software [55] and both combined together to obtain the ratio (Mucilage Area/Seed Area). All of the overlapping released-mucilages were removed from the data set. Data are the means of three biological replicates, each of at least thirty seeds per RILs (90 seeds per RILs).

The oil content and FA composition analysis of the RIL population is described below.

### Characterisation of the Seeds and Embryos

Seed weight was determined on mature seeds with three biological replicates per RILs, each constituted of four seed lots of thirty seeds. Area and circularity of seed and embryo analysis was performed using image segmentation analysis. Segmentation of both tissues outlines from the image background were performed using ImageJ software and the “colour threshold” function [55–57]. Area and circularity measurements were performed on four biological replicates, each of thirty seeds (Supplemental Figure S1).

### Image Analysis

#### Scanning Electron Microscopy (SEM)

Scanning electron microscopy was used to examine the surface of the seed coat and its cellular microstructure. Dry seeds were placed on the mounting stubs and fitted on a Peltier stage and then examined using a high-resolution environmental scanning electron microscope (FEI Quanta 200 FEG). The samples were observed at 2 °C under a variable water vapor pressure using an accelerating voltage of 5 kV. The images were taken at the working distance of 4 mm and varied magnifications of between 800x to 6000x, and saved as TIFF files. The brightness and contrast of the saved images were slightly improved using ImageJ software [55].

#### Starch granules analysis

Analysis of the starch granules content was performed on the toluidine blue O-stained seed sections.

Toluidine blue O staining intensity plots were generated across a transect spanning MSCs from the distal to the basal primary cell walls using ImageJ software [55]. To facilitate the peak detection corresponding to each mucilage layer and because of the purple-pink colour of the mucilage polysaccharides when stained with toluidine blue O, red and blue channels were removed from RGB images.

For the starch granule analysis, images from cytological analysis were processed with ImageJ software; starch granule outlines were manually detected with the freehand selection tool and added to the ROI manager for distinct labelling, i.e. pink for starch in MSCs, blue for starch in parenchymatous cells. Starch granules are composed of several parts and because we are working on thin cytological sections, we could not exclude the risk of detecting fewer starch granules due to their orientation. We thus chose to count each part of the starch granules independently and obtain the mean by the number of cells analyzed.

### Micromechanical Analysis

Mechanical measurements were performed on dry mature seeds. We previously found [20].that the outer cell wall of seeds behaves as an elastoplastic material. Such material has two types of responses to external forces. When a small force is applied and released, the material recovers its initial shape; it is described as elastic. Elasticity is characterized by a quantity known as the elastic modulus: the higher the elastic modulus, the smaller the deformation for a given applied force. When a force exceeding a well-defined threshold is applied, the material is irreversibly deformed and does not recover its shape; it is described as plastic. Plasticity is characterized by a quantity known as hardness: the higher the hardness, the higher the threshold force. We quantified elastic modulus and hardness using a TI950 Triboindenter (Hysitron) coupled to an optical microscope. The optical microscope is used first for acquiring pictures of the seed coat surface and locating radial and distal cell walls of the MSCs. To increase the precision of the micro-indentation measurements on the seed surface, i.e. top of radial cell walls, a shape calibration of the 1-*µ*m diameter conical tip was performed. On each radial and distal cell walls target, a 1000-µN force was applied for 5 sec and released in the same span of time. These parameters are necessary for penetration of the conical tip into the sample, through the primary cell wall, with a depth of 300 nm.

Extraction of the elastic modulus and hardness was based on a classical approach in the mechanics of materials [58]. For this, we computed the slope of the unload-displacement curves (force-indentation retract curve) and the contact area of the conical tip with the seed, using Triboscan and R softwares [59] as previously described [20].

For comparison of the physical properties of the selected RILs, micro-indentation on the radial and distal cell walls were treated as part of the same group and combined together. Two hundred and sixty measurements were performed for each RILs (thirteen seeds per RILs with ten measurements on the radial and distal cell walls).

### Oil Content and FA Composition Analysis

#### Extraction and analysis of the seed oil content

Analysis of the seed oil content was performed on mature dry seeds from all the recombinant inbred lines population harvested in the field in generation F10 and from the three selected RILs harvested in greenhouse after F12 generation.

Seed oil extraction was performed on 100 mg of fresh dried seeds. Seeds were ground in liquid nitrogen. One mL of hexane-isopropanol (2:1) was added to the samples, then they were mixed at 4000 rpm for 5 min. The supernatant was recovered and centrifuged at 8000 rpm for 2 min. The supernatant was collected and ether evaporated.

#### FA composition analysis

Analysis of FA was performed either on the whole seed (for FA content phenotyping of the RIL population) or on the embryo for the three selected RILs during seed development except for 10 DPA (whole seed). Analysis of FA was directly performed on the oil extracts from whole seed (RIL population) or after polar metabolites extraction (for the ^1^H-NRM analysis of the three RILs) on the pellet of the samples. For FA extraction on the embryos of the three RILs, pellets from the ^1^H-NMR analyses were dried using a speedvac for 2 h. Five hundred µl of n-hexane (Fisher) containing 0.005 % (w/v) pentadecane was added to the pellet. Fifty µl of TetraMethyl Ammonium Hydroxide (TMAH) was added to the samples, and the mix incubated for 10 min at 990 rpm at 20 °C. Samples were centrifuged for 10 min at 12,000 rpm and 25 °C. One hundred fifty µl of supernatant was retrieved and transferred in a glass vial. The same extraction procedure was applied on the oil extracts from the whole seeds of the RILs population. Two biological replicates were considered for each RIL of the population for FA content phenotyping and three to seven biological replicates for FA content analysis of the three RILs.

FA methyl esters analysis was performed using a TRACE GC ULTRA gas chromatograph system coupled with a DSQ II quadrupole mass spectrometer (Thermo Scientific, France) and equipped with an automated sample injector. One µl of each sample was injected with a 25:1 split ratio at 240 °C. For this step, the ion source was adjusted to 220 °C and the transfer line to 280 °C. Electron-impact ionization method (EI, 70 eV) was used for the analysis with Helium as the carrier gas, at a flow rate of 1 mL min^-1^. After running the organic phase in a column (lenght: 60 m, inner diameter: 0.25 mm, 0.25 µm liquid membrane thickness, TR-FAME, Thermo Scientific, France) at 150 °C for 1 min, the temperature was increased to 180 °C with a gradient of 10 °C min^-1^ and maintained for 30 s at 180 °C, ramped at 1.5 °C min^-1^ to 220 °C, 30 s at 220 °C, followed by 30 °C min^-1^ to reach 250 °C and maintained at 250 °C for 5 min (run time: 38 min). Each mass spectrum was recorded with a scanning range of 50 to 800 m/z, and X Calibur software (Thermo Scientific) was used for manual peak integration and analysis. FA species were identified by comparing their retention times with those of pure standards (Fame mix C14-C22 Supelco-18917-1AMP). FA and standards were analysed under the same conditions. Pentadecane was applied as the internal standard. FA species concentrations for each sample were normalized against the internal control and expressed as a percentage of the total FA content.

### Transcriptomic Analysis

Once each development stage had been reached, flax seed capsules from the three RILs were cut and opened to collect seeds. Seed coats and endosperms were isolated from embryos using a scalpel, fine forceps and a binocular magnifier. Only seed coats and endosperms tissues have been considered for analyses except for 10 DPA (whole seed). About 200 mg of tissue was collected for each condition and frozen in liquid nitrogen. Total RNA extraction was carried out on ground tissues according to hot phenol purification protocol (modified after [60]). RNA quantification was carried out using a Nanodrop 1000 (Thermo Scientific) and only RNAs with OD ratios 260/280 and 260/230 higher than 2 were conserved. Quality controls were achieved using 2100 Bioanalyzer RNA chips (Agilent Technologies Inc., http://agilent.com), according to the manufacturer’s recommendations. cDNA libraries were generated by reverse-transcription from total RNA (500 ng) using GeneChip WT PLUS Reagent Kit (Affymetrix Inc., Santa Clara, USA), according to the supplied protocol. cDNAs (5.5 μg) were fragmented and labelled using the GeneChip WT Terminal Labeling Kit (Affymetrix Inc., Santa Clara, USA).

Arrays were designed based on the recently Whole-Genome Shotgun (WGS) assembly of flax genome60 resources available on 20-22-mer specific probes covering 43,384 flax genes [61]. Hybridization on the GeneAtlas System (Affymetrix) of the labelled cDNA samples was carried out automatically using the Hybridization Station according to the manufacturer’s recommendations. Hybridized array strips were scanned using the GeneAtlas Imaging Station and GeneAtlas Instrument Control Software (Affymetrix) at 2-μm resolution. After the imaging step, a grid was aligned on the Image (DAT) files to identify the probe cells and compute the spot intensity data to generate Cell Intensity data (CEL) files. All quality control tests for signal value, hybridization, labelling and sample quality were performed using the GeneAtlas software according to the manufacturer’s recommendations. CEL files were exported into the Partek Genomics Suite 6.6 software (Partek) and transformed into CHP Files for gene expression analysis using the Robust Multi-array Analysis (RMA) algorithm for quantile normalization and the general background correction prior to gene expression analysis. Three biological replicates were used for each of the three RILs and for the nine biological conditions considered. The Partek Genomics Suite 6.6 software was used for comparison analysis of the gene expression. A three-way ANOVA (one-factor ANalysis Of VAriance) was applied on the dataset from all oligo microarrays experiments in order to identify differentially expressed genes, assigning the three independent categorical variables to the developmental stages, the tissues and the RILs. Corrections for *p*-values were obtained from multiple tests using the Benjamini and Hochberg equation (0.05 False Discovery Rate threshold) [62]. For pairwise comparison analysis between samples (RIL 80 vs RIL 283; RIL 44 vs RIL 80 and RIL 80 vs RIL 44), several levels of contrast were added in the statistical analysis settings. Significance of the threshold was set as drastic (p<0.05 and –2 < Fold change > 2). The Partek Genomic Suite 6.6 software was used to perform and visualize Principal Component Analysis (PCA) and Venn diagram.

Selection of the gene lists used for expression profile analysis and construction of the histograms corresponding to the enumeration of genes was based on our working hypotheses that varied according to the targeted compounds, i.e. mucilage, FA and GA metabolism, and based on a literature search. The differentially expressed genes detected in the 27 pairwise comparisons between samples (nine experimental conditions with three pairwise comparisons each) were numbered and reported on histograms. Analysis of the expression patterns of the genes (mean of the relative signal intensity – Log2 transformed) involved in the mucilage-, FA- and GA-related metabolisms was carried out after importing of the raw arrays datasets in MATLAB (The Mathworks) and using parallel coordinates plot function for visualisation.

### Metabolomic Analysis

#### NMR extraction procedure

For polar metabolites extraction, we weighed 10 mg (+/- 0.02 mg) of freeze-dried tissues of the whole seed, embryo and SCE of the three RILs and for the nine experimental conditions. Dried tissues were ground into the fine powder using a Mixer Mill grinder (MM 400, Retsch, Germany) with 0.4 mm stainless steel milling balls at room temperature (RT) for 6 min at 30 Hz. Grounds samples then were dissolved in 600 µl of H_2_0/MeOH (*v*/*v*) (UHPLC grade quality). 400 µl of supernatant was retrieved from each sample and then 400 µl of H_2_0/MeOH was added to the remaining pellet. The same procedure was repeated three times and the three supernatants were pooled into a tube vortexed at RT for 5 min, sonicated at 60 °C for 10 min and centrifuged at 4 °C for 25 min at 14,000 rpm. Samples pH was adjusted to 6 (+/- 0.02) using HCl and NaOH.Samples then were concentrated using a speedvac overnight. Pellets of the samples were solubilised in 60 µl of D_2_O/MeOD buffer (D_2_O/MeOD 40/60 (v/v), KH_2_PO_4_ 0,1M, TMSP-*d4* 0,0125 % (w/v) pH6). Samples were vortexed for 10 min, sonicated for 45 min at 60 °C and then centrifuged for 5 min at 4 °C and 14,000 rpm. Fifty µl of supernatant was transferred to mm capillary sample tubes (Bruker, Rheinstetten, Germany). For each of the nine experimental conditions and for the three RILs, 10 to 18 biological replicates were analysed (3 to 6 biological replicates harvested in three different greenhouses).

#### NMR instrumentation and spectra acquisition

NMR spectra measurements were based on the procedure used by [43]. All NMR spectra (^1^H-NMR, 2D ^1^H *J*-resolved, 2D COSY and 2D HSQC) were acquired on a Bruker AVANCE III 600 spectrometer (Magnet system 14.09 T 600 MHz/54 mm) operating for ^1^H at 600.17 MHz and using a multinuclear broadband TXI 1.7 mm z-gradient probe. NMR spectra were recorded at 300 K and using the TOPSPIN 3.2 software (Bruker, Rheinstetten, Germany). Tuning and matching steps of the probes were carried out automatically for each sample. Field/frequency lock and shims were performed on the MeOD, after optimization of the shim with tuning and using Topshim. Optimized parameters were applied to all subsequent samples.

The residual H_2_O signal from δ 4.7 and δ 5 ppm was removed applying a pre-saturation sequence, with low power irradiation at the H_2_O frequency and with a relaxation delay of 7 s. For each sample, 1D NOESY spectra were run with a fixed receiver gain (RG=256), using 128 scans of 256 K data points and with a spectral width of 8403,361 Hz (14 ppm). For improving spectra baseline quality, the FID was multiplied using an exponential weighing function prior to Fourier transformation with a line-broadening factor of 0.3 Hz. 2D 1H JRES (J-resolved) NMR spectra were run with a RG=256, using 16 scans in 128 increments, collected in 32 K data points and with a spectral width of 8403,361 Hz along the chemical shift axis and 60 Hz along the spin-spin coupling axis. 2D COSY spectra were run with a RG=256, using 16 scans in 512 increments and collected in 4 K data points, with a spectral width of 8403,361 Hz for both dimensions. 2D HSQC spectra were run with a RG=2050, using 32 scans in 1024 increments and collected in 1 K data points, with a spectral width of 200 ppm in F1 and 14 ppm in F2.

#### Data analysis

All spectra were manually phased and calibrated at 0.0 ppm to TMSP signal using TopSpin 3.2 software (Bruker, Rheinstetten, Germany). Spectra were exported to ASCII using TopSpin 3.2 software. Spectra were analyzed using Matlab software (The Matworks Inc, Natick, Massachusetts, USA) for baseline correction with package airPLS 2.0 [63]. The data around the region below -0.6 ppm and over 10.3 ppm were removed. In order to facilitate spectra alignment between all experimental conditions, the 1D NOESY spectra for the three RILs were aligned with manual patterns for each stage of development and independently for each tissue (whole seed, seed coat and embryo) with the icoshift algorithm 1.3 [64]. Then, each spectra group corresponding to a specific stage of development was aligned together to obtain final spectra alignment (RILs and stage of development) independently for the three tissues. Finally, patterns of the three tissues were aligned together and the resulting pattern was manually adjusted. The “dynamic adaptive binning” or “DAB” algorithm [65] was applied for automatic bin boundaries detection, spectra bucketing and bins integrations. The resulting data set corresponding to the bins area was analyzed using principal component analysis (PCA) with the SIMCA-P^+^ software (version 12.0.1, Umetrics, Umea, Sweden) applying the Pareto scaling method. Chemical shifts of discriminating signals were identified using loading plots. For identification of the discriminant metabolites for all experimental conditions, a comparison was performed between chemical shifts identified and those of reference compounds (prepared in MeOD/D_2_0, pH 6) and assignments of the discriminant signals was performed using ^1^H spectra, which was also compared with ^1^H spectra of the reference compounds. Results were confirmed by consulting the Biological Magnetic Resonance Data Bank (www.bmrb.wisc.edu) and The Human Metabolome Database (www.hmdb.ca) and using 2D 1H JRES, 2D COSY et 2D HSQC spectra acquired on one representative samples for each of the nine experimental conditions. In addition for some compounds, chemical shifts were compared with previously reported data [43]. HCA visualization of the log-2 transformed data set from resulting area was performed in Matlab. Box plot visualization of the data was performed in R software [59] as well as visualization of the curves describing the sucrose/glucose content evolution.

### Lignan Analysis

Analysis of Lignan profiles was performed on the seed coat tissues previously analysed by ^1^H-NMR. The 1.7 mm capillary samples tubes were broken and the samples were collected, centrifuged at 6,000 rpm for 15 s. Ten µl of supernatant was recovered and deposited on the pellet of the corresponding sample analysed using ^1^H-NMR. Samples were vortexed for 6 min, sonicated for 10 min and centrifuged for 5 min at 14,000 rpm. Ten µl of supernatant was recovered and mixed with 590 µl of H_2_0/MeOH (UHPLC) (*v*/*v*). Samples were hydrolyzed by suspension in 6 µl of 10 M NaOH. After a quick vortex step, samples then were incubated at 60 °C for 3 h with shaking at 2,000 rpm and adjusting their pH with 6 µl of HCl 37 %. Samples were quickly centrifuged at 14,000 rpm at 4 °C and the supernatants were collected and filtered using a Teflon membrane filter with a 0.2 µm pore size. HPLC-UV analysis was performed on 50 µl of each sample using a Shimadzu high-performance liquid chromatograph (LC-2010HT, Shimadzu Scientific Instruments). Separation was performed using a reversed phase column (25 cm L x 10 mm D, 5 µm, Sigma-Aldrich) controlled by the LabSolutions software. The sample was injected with an automatic injector through a 20 µl full loop. Detection of the lignan was performed at 280 nm. For each sample, the chromatogram was reduced to ASCII files and exported in Matlab (R2014b, The Mathworks Inc, Massachusetts, USA). Each Chromatogram was baseline corrected using airPLS algorithm 2.0 [63] and aligned according to the stage of development with icoshift algorithm 1.3 [64] for each RILs, and then they were all combined and aligned together. The peaks corresponding to each of the lignans identified were manual integrated.

### Statistics

For each experiment, normality and homoscedasticity of the dataset was tested using a Shapiro-Wilk and a Bartlett test, respectively and accordingly, a one-way ANOVA test or a nonparametric Kruskal-Wallis *H* test was performed. One-way ANOVA was followed by a Tukey-HSD multiple comparison *post hoc* test and the Kruskal-Wallis by a Mann-Whitney-Wilcoxon *U post hoc* test (α=0.05, after Bonferroni correction). All statistical analyses were performed using Statistica software package (v.9.1, StatSoft, Inc., 1984-2010).

## RESULTS AND DISCUSSION

### Identification and phenotyping of three flax recombinant inbred lines (RILs) displaying contrasted seed oil and released mucilage contents

In order to identify flax lines displaying contrasting phenotype in terms of mucilage release and seed oil contents, a multiple screening analysis was performed on a flax recombinant inbred lines (RILs) population produced by crossing between Oliver (oil-flax) and Viking (fiber-flax) cultivars. Two RILs were selected for severe antagonistic differences, i.e. RIL 283 (low mucilage release, low oil content) and RIL 80 (high mucilage release, high oil content). RIL 44 was chosen as the reference line because of its intermediate phenotypes for both traits, medium mucilage release and oil content (Figure 1a and b, Supplemental Figure S2). The color of the seed coat in flaxseeds varies during seed development before rendering the final color to the seeds (Figure 1c) [66], suggesting the existence of modifications in some seed coat-specific components, including phenolic compounds such as proanthocyanidins [67]. No color difference was found between the selected RILs (Supplemental Figure S3a), suggesting the absence of any major variation in phenolic compounds in mature seed between the 3 lines. It has been shown that the lignan content should be negatively correlated to the ω-3 FA content in flaxseeds [43]. Our HPLC-UV analysis of the lignan content in the seed coat of the three selected RILs, however, has shown no differences (Supplementary Figure S4). Seed size and weight are important targets in determining yields of crop breeding [68,69]. In general, large seeds, which have bigger endosperm and cotyledons, have more advantages in crop yield [70]. The size and the weight of the RIL 80 are higher than those of RIL 44 and RIL 283 (Supplemental Figure S1b, Supplemental Figure S3c). The high weight of RIL 80 can also be explained by the fact that this RIL contains more fatty acids (Figure 1a) and releases more mucilage (Figure 1b, Supplemental Figure S1a) than the RILs 44 and 283.

**Figure 1.**
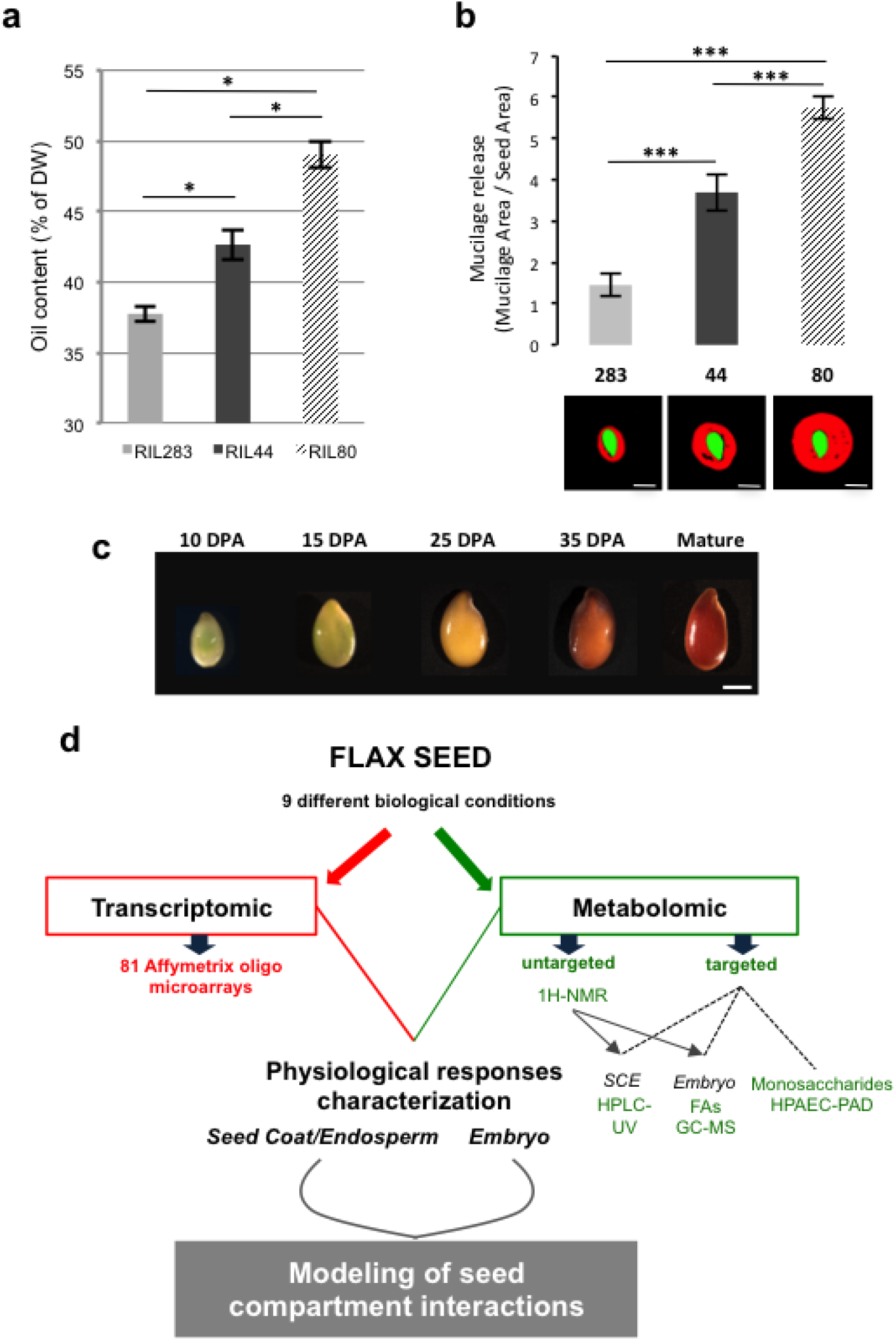
Phenotypic characterization of the 3 selected RILs and experimental strategy overview. a. Quantification of oil content in mature seed from selected RILs. Error bars represent +/- SE (n = 5 biological replicates, each of 100 mg of seeds). Kruskal-Wallis *H*-test followed by a Mann-Whitney *U*-test, \**P*<0.05. b. Assessment of mucilage release in mature seed from selected RILs. Mucilage production was estimated by calculating the ratio between released mucilage area and seed area, both obtained with image analysis using the ImageJ software (see Materials and Methods for more details). Error bars represent +/- SE. Kruskal-Wallis *H*-test followed by a Mann-Whitney *U*-test, \*\*\**P*<0.001. Data are means of three biological replicates. Bars = 4 mm. c. Flax seeds of RIL 44 at different stages of development corresponding to the ones used in this study: from early embryo, i.e. 10 days post-anthesis (DPA), to mature seed, i.e. 60 DPA, showing differences in seed coat color. Bar = 2 mm. d. Overview of the experimental procedure combining transcriptomic and metabolomic analyses performed on each of the 3 selected RILs. A set of 27 experimental conditions were analyzed (i.e. 9 for each of the 3 RILs) consisting of 10 DPA whole seed, 15, 25, 35 and 60 DPA embryo and seed coat/endosperm (SCE).

### Experimental strategy combining transcriptomic and metabolomic analyses during seed development

Transcriptomic and metabolomic analyses have been performed on the three RILs during seed development, at five stages of development flanking the seed oil biosynthesis and the main physiological events linked to the flax mucilage production and modifications [5,20], i.e. 10, 15, 25 and 35 DPA (Days Post-Anthesis) and mature seeds (≈ 60 DPA). For each time point, both analyses have been conducted in parallel on the seed coat/endosperm (SCE) and the embryo tissues. 10 DPA material was analyzed as whole seeds because of the low amount of embryonic tissue and the large quantity of endosperm still present at this stage. Finally, nine biological conditions have been considered for each RILs (Figure 1d). Transcriptomic analysis has been carried out using 81 Affymetrix oligo-microarrays. Metabolomic analysis was performed in two ways: untargeted ^1^H-NMR metabolic profiling performed on every samples and targeted analysis of the mucilage chemical composition which was only performed on mature whole seeds (HPAEC-PAD). Fatty acid (FA) composition of the whole embryo was analyzed using GC-MS after ^1^H-NMR analysis of the polar metabolites (see “Materials and Methods” for more details). The final goal of this work is to model the regulation of embryo and SCE interactions by combining both transcriptomic and metabolomic analyses performed on the phenotypically distinct flax RILs (Figure 1d).

### Partitioning of the main metabolic variations in flax seed tissues during development

Metabolomic profiling techniques have already been used on various crops including model oilseed species, however very little is known about metabolite changes in flax seed during development. The recent development of a ^1^H-NMR metabolomics-based tool enabled the metabolite content analysis of flax seeds [43]. This method is especially well suited for the analysis of amino acids, carbohydrates, carboxylic acids, phenolic compounds and other metabolites involved in the lipid metabolism. This tool has been applied on our selected RILs to investigate metabolic pathways that can be particularly affected between RILs, within tissues and during seed development (Figure 2). Principal component analysis (PCA) performed on the datasets obtained from ^1^H-NMR experiments revealed severe metabolite changes between samples according to the seed tissue and to the developmental stage, as indicated by the first two PCA scores which explained 53 % of the total variability. The first component (PC1) explained 40 % of the total variability and allowed segregating samples according to the kinetic of development, especially for the early stages of development, i.e. 10 to 25 DPA (Figure 2a). The second principal component (PC2) explained 13 % of the total variability and allowed to segregate samples from whole seed (green), SCE (blue) and embryo (red) tissues (Figure 2b). Neither for the SCE nor for the embryo tissues, levels of variability for metabolomes between 35 DPA and mature seed could be clearly assessed from the PCA analysis especially in the embryo (same clusters), which suggests the pause of metabolite changes in the SCE and embryo at around 35 DPA. The PCA analysis also revealed that samples from embryo and SCE followed parallel trajectories according to the horizontal component (PC1), although they displayed clear metabolite differences between tissues and during development. Samples from the whole seed, which contain both tissues and large proportion of non-cellullarized endosperm, appeared closer to the ones from SCE according to PC2. Altogether these results suggest a clear gap between embryo and SCE metabolites profiles especially during the early stages of development. They also suggest that metabolism in early whole seeds is mainly driven by the seed coat rather than embryo. This is in agreement with the fact that biosynthesis of mucilage and some phenolic compounds occurs early in the flax seed coat, and that storage compounds, such as oil and proteins, accumulate later on in the embryo [5,7,20]. Distinguishing samples from the three selected RILs on the PCA scatter plot (Figure 2c) revealed that the RILs did not seem to present significant metabolite changes, with an exception for the whole seed at 10 DPA. Examination of the PCA loading plot allowed the identification of 20 discriminating metabolites, which segregate between RILs in at least one experimental condition (Supplemental Table I). This set of discriminating metabolites has been submitted to a hierarchical cluster analysis (HCA) in order to evidence the level of each metabolite in each seed tissue and developmental stage (Figure 2d). As shown by the dendrogram at the top of the HCA, the log-2 transformed dataset was automatically organized into three major clusters (Clu). The whole seed (10 DPA) and both tissues at 15 DPA clustered together (Clu-1), showing relatively high content of all discriminating metabolites, except for D-(+)-Raffinose and two cyanogenes, i.e. Linustatine and Neolinustatine. For later stages (25 DPA to mature seed), the datasets clustered according to the discriminating metabolites, which especially accumulated in the seed coat (Clu-2) and in the embryo (Clu-3). The D-(+)-Raffinose accumulated in both tissues during development, especially in the embryo, suggesting that the embryo is more sensitive to maturation and desiccation-related events than SCE (Figure 2d). Analysis of the HCA also revealed that the major part of these discriminating metabolites, such as sucrose and glucose, showed a severe relative content decrease in the SCE and the embryo after 15 DPA, confirming the importance of the degradation of these compounds for the seed, allowing production of substitutes and building of new and/or more complex metabolites potentially involved in the mucilage or FA biosynthesis [33].

**Figure 2.**
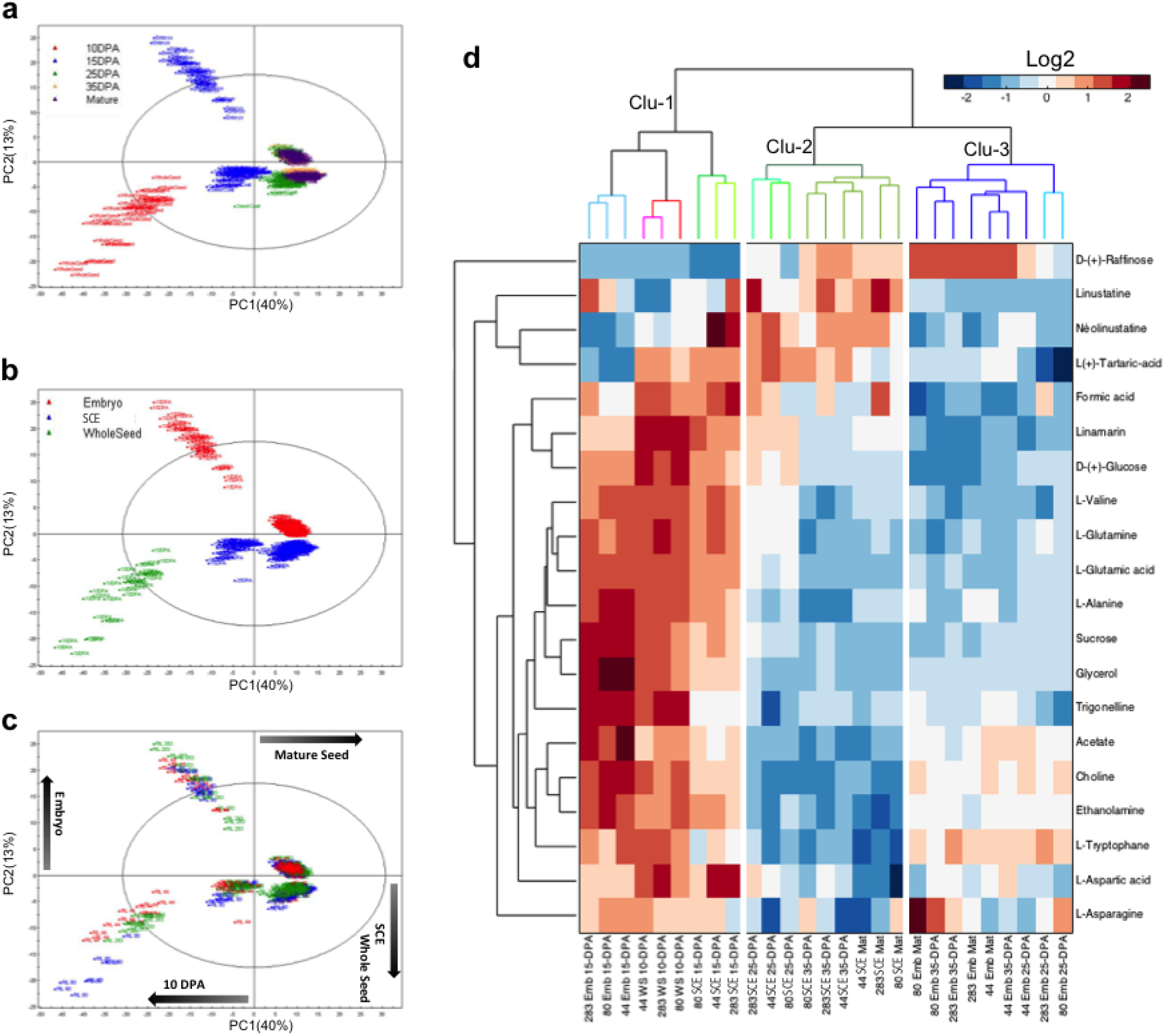
Metabolite profiling in flax RILs during seed development. (a-c) PCA scores plot of the 1H-NMR experiment performed on the three selected RILs during seed development and according to the specific tissues analysed. PCAs are labelled according to the developmental stages following the horizontal component (a), tissues following the vertical component (b) or the selected RILs (c). (d) HCA of the discriminant metabolites identified from metabolite profiling according to the 27 different experimental conditions. The colour scale represents mean intensities (Log-2 transformed) of the relative metabolite content. Data are means of 10 to 18 biological replicates. The dendrogram colour code illustrates hierarchical clustering according to the tissues (10 DPA Whole seed, red; Embryo, blue; seed coat/endosperm (SCE), green). Clu, cluster.

As suggested by PCA and HCA analyses and the few numbers of identified discriminating metabolites, metabolome rearrangements in the embryo and SCE look similar between RILs during seed development and do not seem to sustain the differences in mucilage and seed oil contents observed in these lines, except for sucrose, glucose and glycerol since they are key entries of the sugar and TAG metabolism. Thus, phenotypical differences observed between the RILs seeds are not likely to come from major modifications in the primary and secondary metabolisms. These differences could be due to changes in other classes of metabolites such as non-polar volatiles or lipids compounds including FAs, justifying our analysis of the FAs content and some other seed coat specific compounds using targeted metabolomics.

### The seed coat is a key compartment for transcriptional regulations occurring at midcourse of seed development

Previous transcriptomic analyses on flax seed have identified the seed coat as a specific site of transcriptional regulations occurring during seed development and the tissues where lignans, flavonoids, and mucilage biosynthetic pathways take place [25,66,71]. To determine if the mucilage and oil content variations found in the selected RILs could be sustained by RILs-specific transcriptional regulations, a transcriptomic analysis using flax Affymetrix oligo-microarrays was performed on the same set of samples as for the metabolomics analysis (9 samples for each line; Figure 1d). PCA of the dataset revealed clear transcriptome changes between samples (Figure 3a-b), PCA1 explaining 28.7 % of the total variability and segregating samples according to the development stages (Figure 3b), and PCA2 explaining 8.6 % of the total variability and segregating samples according to the specific tissues (Figure 3a). A three-way ANOVA analysis was performed on the oligo-microarrays dataset from all experimental conditions followed by a pairwise comparisons analysis to identify the number of genes differentially expressed between two RILs according to the tissues and to the developmental stages. RIL 283 produces low oil/mucilage contents, whereas RIL 80 produces high contents and RIL 44 displays intermediate phenotype. Therefore, in order to identify putative correlations between phenotype intensity and specific gene expressions, genes differentially expressed in SCE and embryo between 2 of the 3 RILs were identified using the transcriptomic data obtained, comparing RIL 44 vs RIL 283 and RIL 80 vs RIL 283 and RIL 80 vs RIL 44. The Venn diagrams from Figure 3c and d were obtained by combining the 3 comparisons. It clearly shows that the higher gene expression regulations (up- or downregulations) occur in the SCE at 25 DPA. At this stage of flax seed development, the number of genes which are differentially expressed is high in RIL 80 vs RIL 283 (1381 up; 450 down) and in RIL 44 vs 283 (993 up; 351 down), only a few genes seem to be differentially expressed in RIL 80 vs RIL 44 (45 up; 19 down). This can be explained by the fact that the lists consist mainly of genes with slight differential expression, making a ratio significant (just above the threshold of significance, i.e. the gene is in the list) in one comparison (i.e. in RIL 80 compared to RIL 283) while the ratios will be just below the threshold (i.e. the gene is not in the other lists) in the other comparisons (i.e. in RIL 44 compared to RIL 283 and in RIL 80 compared to RIL 44). In the embryo, the highest number of differentially expressed genes is also found at 25 DPA in RIL 80 vs 283 (442 up; 175 down) and RIL 44 vs 283 (134 up; 99 down). Very few genes are differentially expressed between RIL 80 and 44 in this tissue at 25 DPA.

**Figure 3.**
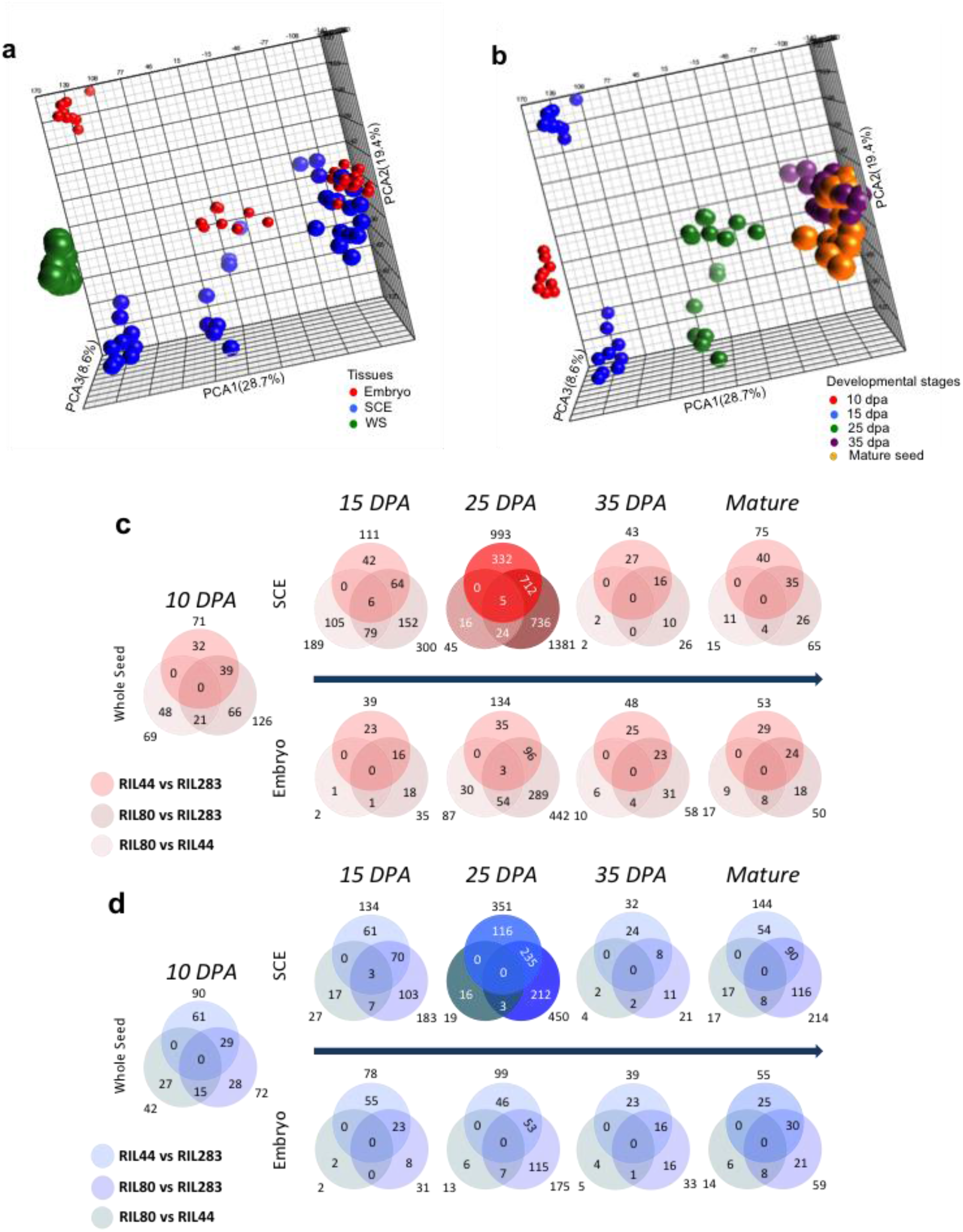
Transcriptomic analysis in flax RILs during seed development. (a,b) 3D-PCA scores plot of the dataset from the 81 Affymetrix oligo microarrays analysis performed on the three selected RILs. PCAs are labelled according to the specific tissues following the vertical component (a) and the developmental stages following the horizontal component (b). Various sizes were applied to each sample attribute for a better display of the results. (c-d) Pairwise comparisons analysis of the number of genes differentially expressed between the three selected RILs. Venn diagram show the number of unique and overlapping genes upregulated (c) or downregulated (d) between at least two RILs. A three-way ANOVA (Fold-change > 2 (c) or < -2 (d), *p*-value adjusted with FDR < 0.05) was used for extraction and combination of the genes lists. Bold circles represent the most important lists of unique and overlapping differentially expressed genes.

Thus, the transcriptomic analysis clearly revealed great differences in the transcriptomes of the three RILs, especially well pronounced in the SCE at 25 DPA and mainly overexpressed in RIL 80 and 44 vs RIL 283. Despite no major differences have been observed between RIL 44 vs 80, we hypothesized that phenotypic variations between RILs for the released mucilage and seed oil contents could be mainly driven by the SCE in the middle of the seed development at the transcriptomic level.

### A large set of genes related to cell wall, fatty acid and gibberellin is differentially expressed in 25 DPA seed coat/endosperm

In order to gain insight into the transcriptional regulations observed in the SCE at 25 DPA, we identified, among the genes differentially expressed in RIL44 vs RIL283, RIL80 vs RIL283 and RIL80 vs RIL44 comparisons, the ones for which the level of up- or down-expression is ≥ 2 (Supplemental Figure S5). The number of up- or down-regulated genes has been selected on an arbitrary basis. A larger number of genes are up or down-regulated in the SCE of RIL 44 vs RIL 283 (1344 genes) and in the SCE of RIL 80 vs 283 (1831 genes) compared to those regulated in the SCE of RIL 80 vs 44 (only 64 genes). Families of up or down-regulated identified genes are the same in SCE for RIL 80 vs RIL 283 and RIL 44 vs RIL 283, with higher expression level in RIL 80 vs RIL 283.Among the genes up-regulated in RILL 44 vs RIL 283, genes putatively involved in cell wall metabolism, in mucilage biosynthesis or modifications (peroxidases, beta-galactosidases and beta-glucosidases, fasciclin-like arabinogalactan proteins (AGP), UDP-glycosyltransferases, glycosyl hydrolases, pectin lyase-like proteins) are found. Genes putatively involved in the seed oil metabolism, essentially in poly-unsaturated fatty acid and very long chain fatty acid biosynthesis (lipid transfer proteins, long-chain alcohol dehydrogenase, fatty acid desaturase 3 and 7 (FAD) and plant stearoyl-acyl-carrier-protein desaturase protein (SAD)) are also up-regulated in the SCE of RIL 44 vs RIL 283 and RIL 80 vs 283. Genes putatively involved in the gibberellic acid pathways like oxidases, GA20ox and GA3ox, GA-regulated family proteins and GA requiring 3 were up or down-regulated in the SCE of these RILs (Supplemental Figure S5). Gibberellins are phytohormones that are very well-known for their roles in the control of seed germination, seed weight and size, desiccation tolerance, seed coat formation but also on seed storage fatty acid and mucilage production [51,52,54,72,73]. At last, the number of genes down-regulated in the SCE of RILs was lower than the number of genes up-regulated (Supplemental Figure S5).

### Differences in stage-specific seed coat transcriptome regulations correlate with the differences in mechanochemical regulations of the mucilage release observed between RILs

Differentiation of the MSCs in Arabidopsis involves a complex regulatory network based on a hierarchical cascade model of master regulatory genes, transcription factors and some of their target genes [2,74,75]. Very little is known about the transcriptional regulation of MSC’s differentiation in other agronomic oilseed species, like flax [20]. To determine whether the differences found in the mucilage-related processes between RILs are linked to spatio-temporal differences at the transcriptional level, we numbered the genes potentially involved in mucilage and cell wall compounds-related regulatory processes which were found differentially expressed in transcriptomic data from Figure 3 (Supplemental Figure S6). Each gene that was differentially expressed between the RILs in at least one of the 27 experimental conditions was considered. The highest number of differentially expressed genes is found in the whole seed and the SCE between 15 to 25 DPA, corresponding to the stages of mucilage biosynthesis and modifications in flax [20]. Among the lists of pairwise differentially expressed genes (Figure 3c and d, Supplementary Figure S5) we focused on those potentially involved in the biosynthesis and modifications of flax mucilage polysaccharides and in mucilage release. Because RILs 44 and 283 did not show differences in their soluble mucilage composition and sensitivity to water hydration [20], we have selected the genes differentially expressed in the seed coat from 10 to 25 DPA between RIL 80 vs RILs 44 and RIL 80 vs 283 but being unaffected in RIL 44 vs RIL 283. Among these genes, only those specifically expressed in at least one of these two development stages were selected to reduce the number of candidates to analyze, excluding all the genes expressed at any other stage. Sixteen categories of mucilage-related genes have been considered (Figure 4). Genes corresponding to putative galacturonosyltransferases and glycosyl hydrolase family proteins are the most representative differentially expressed gene families. The sixteen categories have been clustered based on an exhaustive literature search according to their functions on the biosynthesis and modifications of the flax mucilage polysaccharides, mucilage release and cell wall. Means (Log-2 transformed) of the gene expression values were plotted along seed development (Figure 4). The six parallel coordinate’s plots revealed that main differential transcriptional regulations occurred from 10 (whole seed) to 25 DPA.

**Figure 4.**
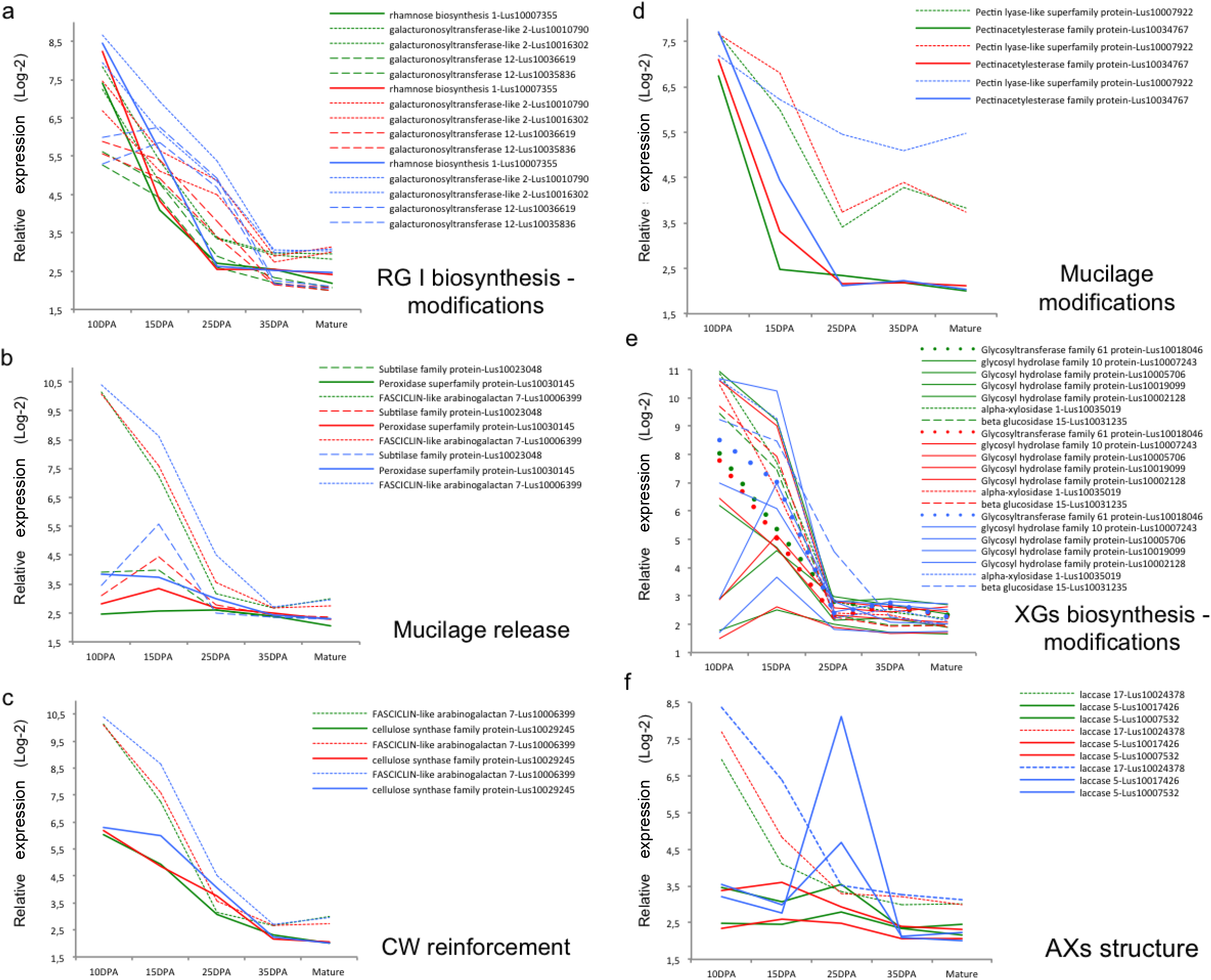
Expression patterns of the genes involved in the chemical composition of the mucilage and related processes. Genes are selected as especially differentially expressed between RIL 44 and 283 vs RIL 80 in whole seed (10 DPA) and seed coat tissues. Selected genes are classified in six different categories according to their putative roles based on literature. For each one, the corresponding parallel coordinates plot displays the relative gene signal intensity according to the stages of development. The relative signal intensity represents the lsmean (Log-2 transformed) of the raw data extracted from the oligo microarrays analysis. Data are lsmeans of three biological replicates. Each of the genes is associated with its corresponding gene-ID. The FASCICLIN-like AGP 7 was included in two categories according to the literature. RIL 283 is shown in green, RIL 44 in red and RIL 80 in blue. The format of the lines in each plot was chosen to improve data visualization. RG I: Rhamnogalacturans; CW: Cell Wall; XGs: Xyloglucans; AX: Arabinoxylans.

Regarding RG I, putative rhamnose biosynthesis, galacturonosyltransferases (GAUT) and galacturonosyltransferase-like (GAUT-like) were found as specifically overexpressed in the seed coat in RIL 80 when compared to the two other RILs (Figure 4a). It has been shown in flax seed that RG I are synthesized first, followed by XGs (Xyloglucans) and by a presumed AXs synthesis and cell wall reinforcement from 15 to 25 DPA [20]. To identify which of the six clusters of genes could explain phenotypic differences in RIL 80 vs to others RILs, stage-by-stage comparison analyses of the differential transcriptional activities have been performed. At 10 DPA, a putative rhamnose biosynthesis 1 gene (Lus10007355) was found as overexpressed in RIL 80 and RIL 44 vs RIL 283 (Figure 4a). Lus10007355 was also found as overexpressed in the RIL 80 vs the other RILs at 15 DPA. In *Arabidopsis thaliana*, it has been suggested that UDP-glucose could be converted into UDP-rhamnose to contribute to the RG I synthesis by an UDP-Rha synthase 2 (MUM4/RHM2) [76–78]. Thus, the differential regulation of the rhamnose synthase 1 does not correlate with the differences observed in the chemical composition of the soluble mucilage between RILs. This shows that function of the rhamnose synthases would not be maintained and would vary between species, explaining partially diversity between flax and Arabidopsis mucilage regulatory networks as previously observed [66]. At 15 DPA, number of the genes differentially expressed in RIL 80 were found as putative orthologs of genes involved in RG I and XGs synthesis and modifications, cell wall (CW) reinforcement and mucilage release. This list included two putative GAUT and two putative GAUT-like genes. Functional analysis of the two putative GAUT and GAUT-like (AtGATL5 and GAUT11) in Arabidopsis revealed that the mucilage of the corresponding mutants contained reduced galacturonic acid and rhamnose levels [79] and showed decreased levels of soluble mucilage content for GAUT11 [80]. Although the lists of differentially expressed genes do not include the putative orthologs of Arabidopsis GATL5 and GAUT11, the overexpression of the putative GAUT12 and GAUT-like2 in RIL 80 at 15 DPA shows the same differential expression as found for the putative rhamnose biosynthesis 1. One possible explanation could match with a reduced conversion of glucose to rhamnose and galacturonic acid polymerization in RIL 80 during early stage of seed development (before 10 DPA), that would be counterbalanced by the increase of transcriptional activities of rhamnose synthase, GAUT and GAUT-like genes without leading to a chemical phenotype compensation. Transcriptional activity of these genes in RIL 80 could also act on the cell wall synthesis of other seed coat cell layers. At 25 DPA, most of the genes overexpressed in RIL 80 were found as putative orthologs of genes involved in the mucilage release and modifications and in the CW reinforcement (Figure 4b, c and d). For RG I biosynthesis and modifications-related genes, at 25 DPA, the putative GAUT-like2 and GAUT12 expression levels are close between RILs 44 and 80 and overexpressed when compared with RIL 283 (Figure 4a). Thus, differences observed between RILs 44 and 283 for their sensitivity to water hydration could be due to the overexpression of GAUT and GAUT-like genes in RIL 44 especially at 25 DPA.

Two putative laccases were also found as differentially expressed in RIL 80 vs the two others with very high fold-changes (Supplemental Figure S5). Interestingly, laccase can increase arabinoxylans network and their extractability from wheat flour water-extractable arabinoxylan (WEAX), through cross-linking proteins to arabinoxylans [81,82]. The flax mucilage is also mainly composed of arabinoxylans [46,83–86]. Two putative laccases, i.e. laccase 17 and laccase 5, were found as overexpressed in RIL 80 respectively at 15 DPA and at 25 DPA (Figure 4f). These results suggest that the high sensitivity of the RIL 80 to water hydration should not only be linked to its high AXG/RG I ratio but also to a more complex AXs network. Altogether, these results show that differences in the mucilage chemical composition and sensitivity to water hydration between RIL 80 and the others could originate from temporal-specific differential activities of miscellaneous classes of proteins and enzymes. These proteins and enzymes would mainly be involved in RG I and XGs synthesis/modifications from 10 to 15 DPA, and in the mucilage release, CW reinforcement, mucilage modifications and probable modifications of the AXs network from 15 to 25 DPA.

### Differential transcriptional regulation of the seed oil metabolism originates in part from the seed coat

A previous study on Arabidopsis has proposed that disrupting rhamnose synthesis in the seed coat leads to an increase of the seed oil content [33]. Since a rhamnose synthase but also putative GAUT and GAUT-like are differentially expressed between RILs, a focus was made on the seed oil and FA transcriptional regulation. Moreover, pairwise comparisons of the transcriptomic datasets at 25 DPA in the seed coat identified genes with high fold-changes which were putative orthologs of genes involved in the seed oil and FA metabolism (Supplemental Figure S5). To determine whether differences in the oil content profiles between RILs could be linked to spatio-temporal differences at the transcriptional level, genes differentially expressed between RILs (Figure 5) were clustered in sixteen categories and then numbered. Results showed that the higher numbers of differentially expressed unigenes were found in the seed coat at 25 DPA, most of them being overexpressed in RIL 80 vs 283 and in RIL 44 vs 283. A high number of differentially expressed genes was also found in the embryo at 25 DPA but only between RIL 80 and 283, which might be related to the difference in seed oil content found in these RILs. Thus, phenotypic differences for seed oil content accumulation between RIL 283 vs the other could be driven by the transcriptomic level in the seed coat at 25 DPA.

**Figure 5.**
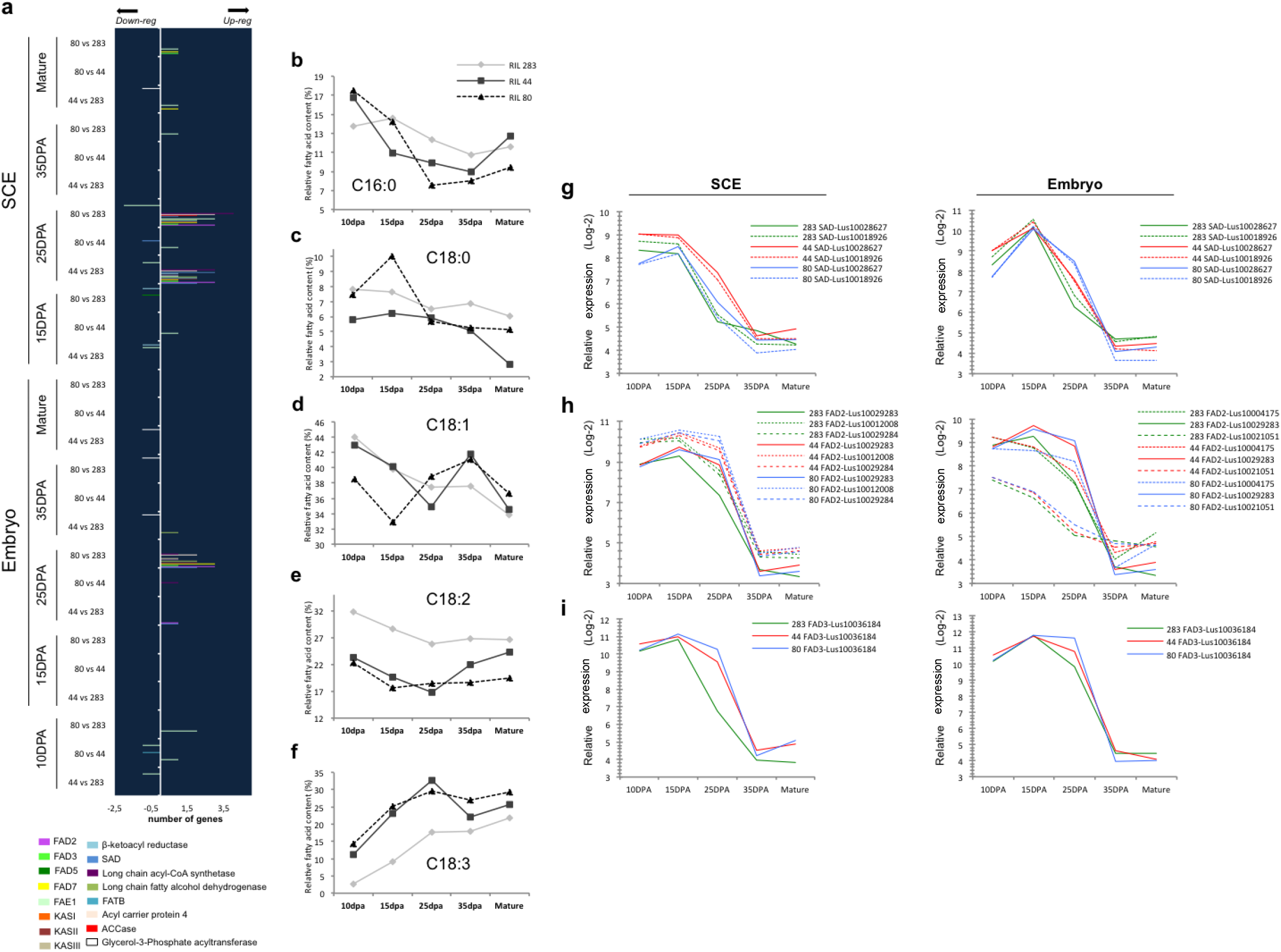
Analysis of the transcriptional regulations of the FA and seed oil metabolisms in flax RILs during seed development. (a) Histogram showing the number of unique and overlapping differentially expressed genes from pairwise comparison analysis of the three RILs. Genes are classified in sixteen different categories according to their putative roles in FA and seed oil metabolism based on literature. (b-f) FA accumulation profiles in the embryo of the three RILs. (g-i) Expression patterns of the genes involved in FA and seed oil metabolism and corresponding to the main fatty acid desaturases; i.e. SAD (g), FAD2 (h) and FAD3 (i). To facilitate data visualization, the expression patterns were displayed separately for the SCE (left) and the embryo (right). Expression patterns of genes of RIL 283, RIL 44 and RIL 80 are respectively represented in green, red and blue. For each one, the corresponding parallel coordinates plot displays the relative gene signal intensity according to the stages of development. The relative signal intensity represents the lsmean (Log-2 transformed) of the raw data extracted from the oligo microarrays analysis. Data are lsmeans of three biological replicates. Each of the genes are associated with their corresponding gene-ID.

In order to determine whether a correlation could be made between the transcriptional regulations in the seed coat and the FA metabolism in the embryo, we characterized the kinetics of FA accumulation in the different RILs and compared them to the microarrays dataset. Results revealed distinct patterns of FA accumulation between the RILs during embryo development (Figure 5b-f). The most interesting observation was that RIL 283 showed different C18:2 and C18:3 contents from 10 to 25 DPA when compared to RIL 44 and RIL 80 (Figure 5e, f). Because fatty acids desaturases act sequentially and play a key role on the flax seed alpha-linolenic acid (ALA, C18:3) accumulation [87], a focus was done on the expression of putative flax orthologs of the main putative fatty acid desaturases including SAD, FAD2 and FAD3 (Figure 5g-i). Two putative orthologs of Arabidopsis SAD, three FAD2 and a FAD3 were found as differentially expressed between RILs during development. Analysis of the parallel coordinated plots revealed high relative expression of the three classes of desaturases from 10 to 25 DPA in embryo and SCE tissues (Figure 5g-i). In the embryo, one SAD is overexpressed in RILs 44 and 80 vs RIL 283 at 25 DPA (Lus10028627). In the seed coat, however, both SAD (Lus10028627 and Lus10018926) are overexpressed in RIL 44 vs the two others at 25 DPA. SAD desaturates C18:0 FA (stearoyl-ACP) to produce C18:1 FA (oleic acid). These results suggest that the low stearic acid content in the embryo of RIL 44 could originate from the overexpression of the SAD in the seed coat at 25 DPA.

For both putative flax orthologs of FAD2 and FAD3, analysis of the relative expression profiles revealed that the flax genes were overexpressed in RIL 80 and 44 vs 283, mainly at 25 DPA in both tissues. The main differential expressions were found in the seed coat (Figure 5g-i). Although all FAD2 and FAD3 were downregulated in RIL 283 compared to the two other lines, comparisons of the differential transcriptional levels with the C18:2 and C18:3 FA accumulations in the embryo did not make sense. However, a correlation has been previously shown between the transcriptional activity of these three desaturases and the accumulation of alpha-Linolenic acid (ALA) [87]. We propose that the differences found between RILs in expression of flax putative orthologs of both FAD2 and FAD3 were sequentially compensated, resulting in a similar accumulation of C18:2 and C18:3. Thus, accumulation of C18:2 and reduction of C18:3 in RIL 44 could originate from the differential expression of SAD, in the seed coat, in the mid-stage of seed development. Therefore, the transcriptional regulation of the seed oil and FA metabolisms seems to originate, at least in part, from the SCE and occurs at 25 DPA during the mucilage modifications step, once the mucilage synthesis is established.

### Mucilage-related phenotypes in RILs are correlated with glucose and sucrose contents in the seed coat up to mid-stage of seed development

It has been proposed recently that seed oil and mucilage biosynthesis shared the same carbon source from the conversion of sucrose into glucose and fructose [33]. The sucrose, which is not used in mucilage synthesis, seems to be loaded into the endosperm to finally reach the embryo via apoplastic connections, contributing to the seed oil production at the very early stages of seed development [33,47]. The first developmental stage analyzed in this study was 10 DPA, which corresponds to early torpedo stage in flax [66] (Supplemental Figure S3b) when no more symplastic and apoplastic connections are supposed to be available between embryo and SCE, precluding any sucrose flux within these seed compartments [47,88]. In order to gain insight into the potential role of sucrose and glucose in the phenotypic differences observed in the RILs seeds, a focus was applied on data related to sucrose and glucose contents (Figure 6). The results showed that sucrose quantities are the same in SCE from the 3 RILs from 10 to 15 DPA, while glucose contents are the same in RILs 44 and 283 but higher in RIL 80 (Figure 6a). From 25 DPA to mature seed, however, glucose contents seem to be the same in RIL 44 and 80 but lower in RIL 283. On the contrary, in the embryo sucrose and glucose levels are very close in the three RILs during seed development (Figure 6b). These results fit well, for the seed coat, to the chemical composition and the soluble mucilage content phenotypes observed. RILs 44 and 283 seem to use the same glucose content from 10 to 15 DPA, during the mucilage synthesis, and they display the same mucilage polysaccharides content with more RG I and less AXGs polysaccharides (Arabinoxylans and Xyloglucans) [20]. The results also suggest that the differential use of sucrose and glucose in the seed coat is not correlated to the levels of each sugar in the embryo. This is in accordance with the hypothesis that from 10 DPA to mature seed, no major apoplastic and symplastic connections enable carbon allocations from the SCE to the embryo.

**Figure 6.**
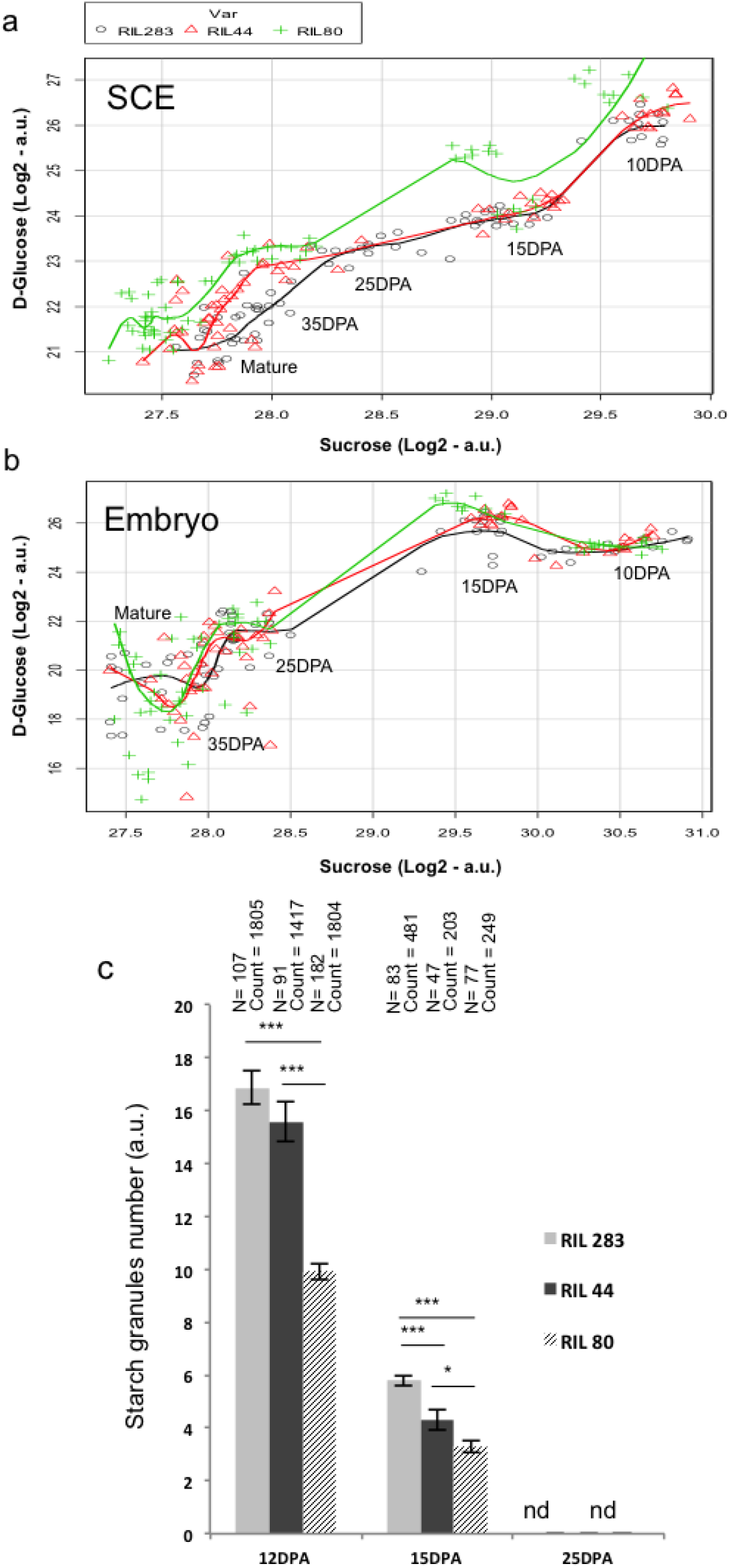
Analysis of the differential use of the main carbon sources in flax RILs during development. (a) Evolution of the sucrose and glucose contents in the SCE. Data are from 1H-NMR experiments. Values are means (Log2 transformed) calculated with 10 to 18 biological replicates. (b) Evolution of the sucrose and glucose contents in the embryo. Data are from 1H-NMR experiments. Values are means (Log2 transformed) calculated with 10 to 18 biological replicates. (c) Starch granules count in the seed coat mucilage secretory cells. Kruskal-Wallis *H*-test followed by a Mann-Whitney *U*-test, \**P*<0.05, \*\*\**P*<0.001. Starch granules count is based on segmentation image analysis using ImageJ software. N, number of cells analysed. and, not detected.

### Mucilage-related phenotypes in RILs are correlated with kinetic of starch granules degradation in the MSCs up to mid-stage of seed development

Starch granules depletion in the seed coat MSCs has been associated to columella shape reinforcement in Arabidopsis [89–91]. In flax, starch granules depletion is concomitant to the sequential synthesis of various MSC layers occurring from 10 to 25 DPA, where the mucilage polysaccharides are mainly synthesized [20]. In addition, the release of flax mucilage is linked to its chemical composition [20]. By counting the starch granules in MSCs using segmentation image analysis (Figure 6c), it appears that RILs 283 and 44 contain more starch granules at 12 DPA compared with RIL 80 and degrade more starch granules compared with RIL 80 from 12 to 15 DPA, during mucilage synthesis. Thus, the similar chemical composition of the mucilage for the RILs 44 and 283 [20] could in part originate from their similar use of the starch granules from 12 to 15 DPA. However, RIL 44 releases more mucilage than RIL 283 and less than RIL This phenotype could be explained by the fact that RIL 44 uses more starch granules from 15 to 25 DPA compared with RIL 283, modifying in turn its mucilage content and affecting its properties, and thus getting closer to the phenotype of the RIL 80 for the soluble mucilage content.

Thus, it is proposed that between 10/12 DPA to 15 DPA, the glucose from the starch is used in the same way for RILs 44 and 283 in the SCE, corresponding to the RG I and AXs/XGs biosynthesis steps. Between 15 to 25 DPA, RIL 44 gets closer to RIL 80 for starch granules degradation, likely for the modifications of the mucilage chemical composition and for increasing/modifying its soluble mucilage content.

### GA-related genes are differentially expressed in RILs seeds

GA metabolism in Arabidopsis seeds has been reported to be involved in the seed coat formation through the degradation of starch granules [51], in the mucilage release [51,52], and in the transcriptional regulation of cell wall–associated proteins [53,54]. GA metabolism can also act upstream of the fatty acids metabolism [42]. Interestingly, in the seed coat, the expression of some GA-related genes was found as differentially <u>expressed</u> between RILs at 25 DPA (Supplemental Figure S5). In order to focus on the regulation of GA-related genes during flax seed development, we extracted the data from our transcriptomic analysis (Figure 3) for flax genes identified as putative orthologs of Arabidopsis genes belonging to the GA metabolism-related genes (i.e. GA3ox, GA20ox). The Figure 7 shows the expression kinetic of some of these flax genes during seed development in the 3 RILs. While each flax gene displayed a specific expression kinetic during the whole developmental process, at 10 and 25 DPA some variations occurred in expression regulations between the RILs. The expression of putative GA3ox3 flax ortholog (Lus10013134) and putative GA20ox4 flax ortholog (Lus10006722) are lower in RIL 283 compared to the 2 other RILs in the SCE from 10 to 15 DPA and at 25 DPA, respectively (Figure 7). In addition, the expression of putative GA3ox1 flax ortholog (Lus10023113) is lower in the SCE from RIL 44 at 25 DPA compared to the 2 other RILs. In the embryo, Lus10013134 also revealed a different expression pattern between the three RILs, downregulated in RIL 283 vs the others at 10 DPA and 25 DPA and upregulated in RIL 44 vs the others at 15 DPA. On the contrary, expression profiles of Lus10006722 and Lus10023113 in the embryo are quite similar between the 3 RILs.

**Figure 7.**
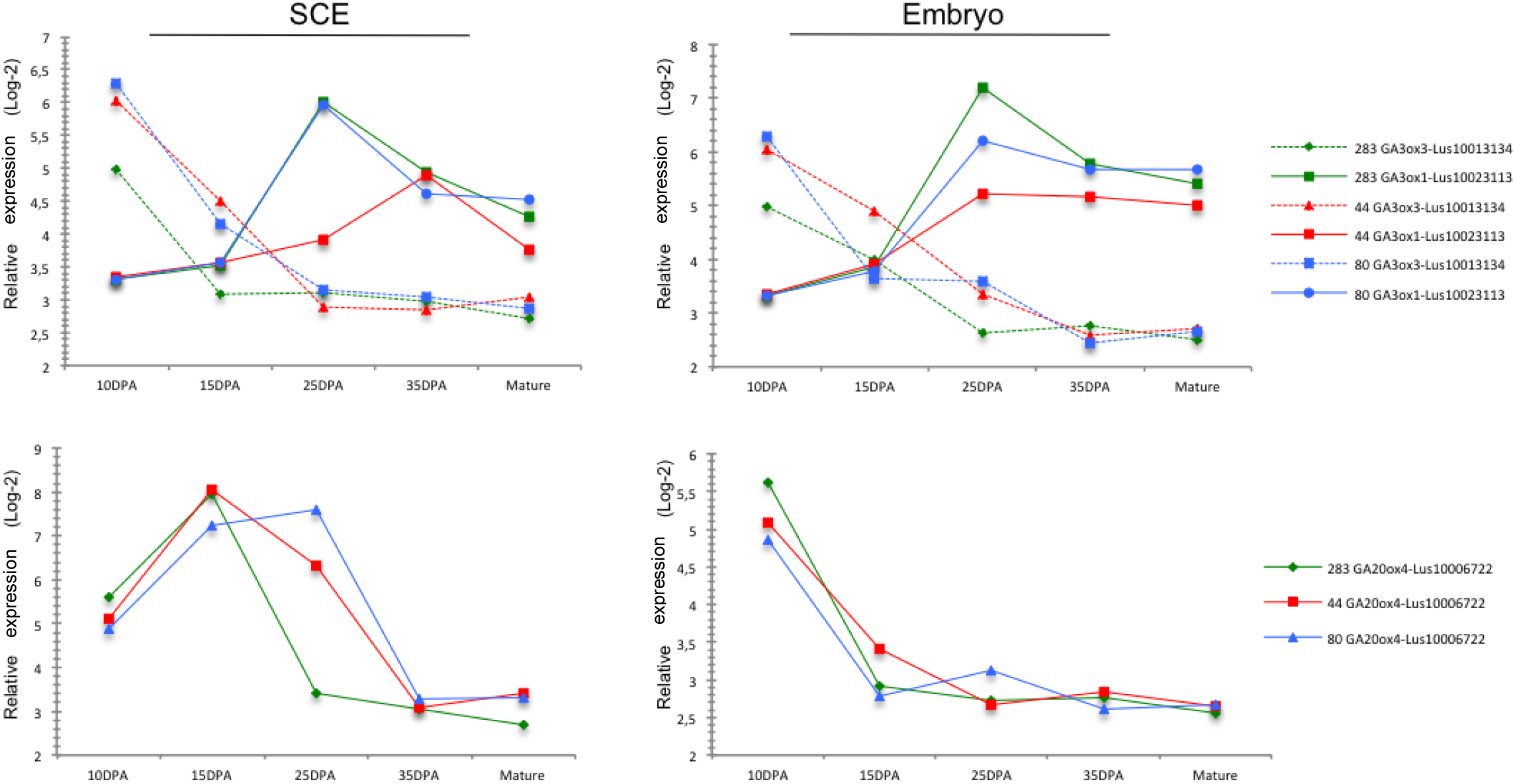
Expression pattern of significantly changes genes involved in the GA metabolism in flax RILs during seed development. For GA3ox or GA20ox genes, the corresponding parallel coordinates plot displays the relative genes signal intensity according to the stages of development. The relative signal intensity represents the lsmean (Log-2 transformed) of the raw data extracted from the oligo microarrays analysis. Data are lsmeans of three biological replicates. Each of the genes are associated with their corresponding gene-ID. To facilitate data visualisation, the expression patterns were displayed separately for the SCE (left) and the embryo (right). RIl 283 is labelled in green, RIl 44 in red and RIL 80 in blue.

### Seed shape and physical analyses suggest a mechanical induction of seed growth and gene expression regulation in seed coat

In this work, we have shown the existence of differences in transcriptional regulations between RILs occurring between 10 to 25 DPA SCE, mainly in 25 DPA SCE, which affect a set of putative flax orthologs of genes involved in mucilage, seed oil and GA metabolism. It has been recently shown that mechanical forces from the endosperm and embryo could be perceived by the seed coat, probably early in seed development, which induces, in turn, a mechano-sensitive regulation of the expression of gene involved in the seed growth and GA metabolism, *ELA1* [50]. Such a similar mechano-sensitive regulation could trigger in flax seed, through GA metabolism modifications and the differential transcriptional regulations observed in our RILs for genes putatively involved in mucilage and FAs/Oil pathways. Image segmentation analysis was performed on mature seeds from the three RILs (Figure 8c) to determine the shape parameters of seed and embryo (Figure 8a-e). Results show that the seed area of the RIL 283 is smaller than for the 2 other RILs, without showing differences between RIL 44 and 80 (Figure 8a). On the contrary, RIL 80 displays a smaller circularity compared to the 2 other RILs, without any differences between RIL 283 and 44 (Figure 8b). RIL 44 also shows an intermediate embryo area (Figure 8c). With respect to the embryo circularity, RIL 44 and 80 are rounder than RIL 283, without any differences among the 2 latter (Figure 8d). Thus, these results indicate that the RIL 44 is close to the RIL 283 for its circularity parameter, while being closer to RIL 80 for the seed size and the embryo circularity, and showing intermediate phenotype for the embryo size.

**Figure 8.**
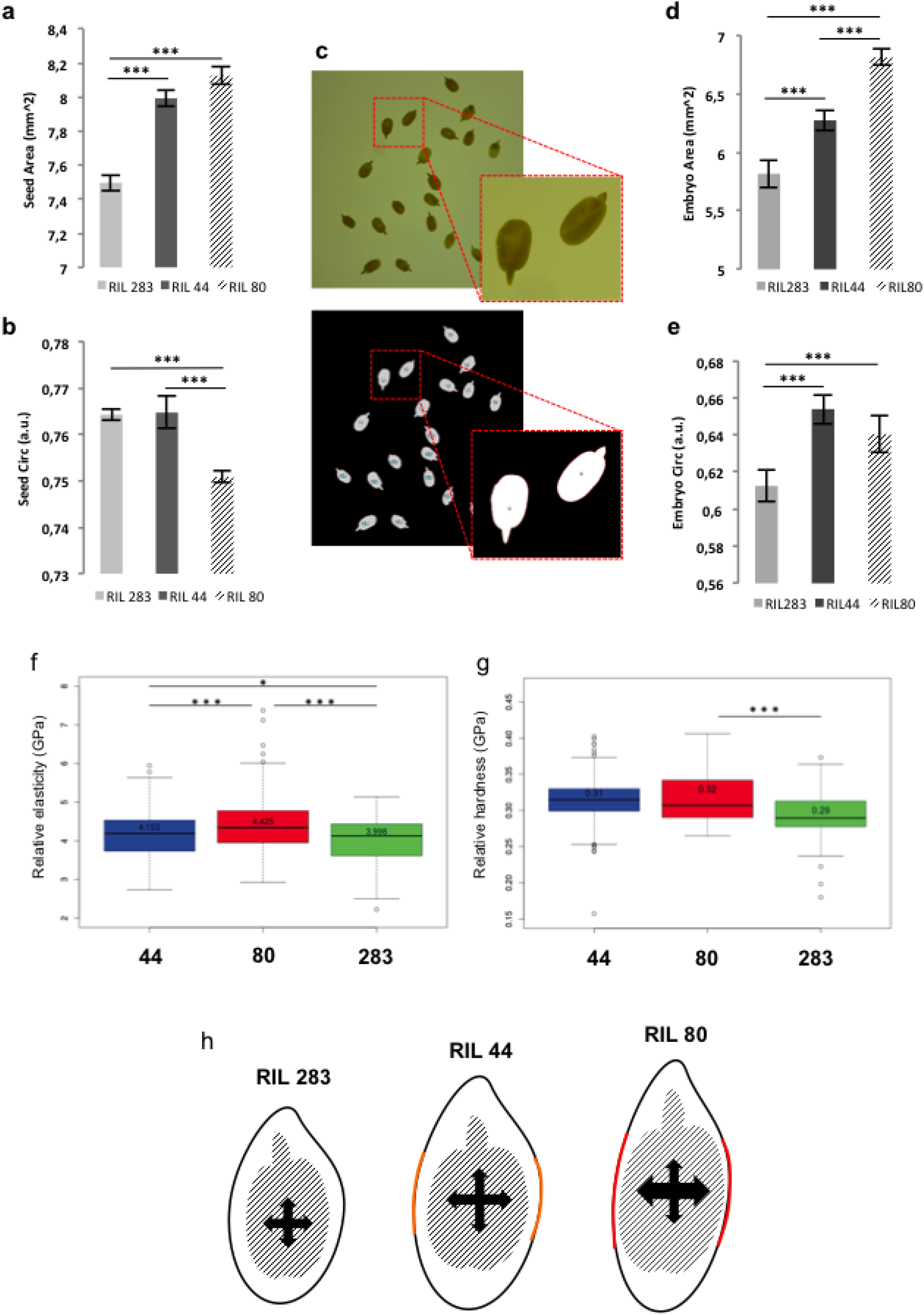
Analysis of the seed growth control through mechanical forces. (a) Seed area measurements on mature flax seeds. Values are means of four biological replicates, each of thirty seeds. Error bars represent +/- SE. Kruskal-Wallis *H*-test followed by a Mann-Whitney *U*-test, \*\*\**P*<0.001. Area measurements was performed using segmentation image analysis using ImageJ software. (b) Seed circularity measurements on mature flax seeds. Values are means of four biological replicates, each of thirty seeds. Error bars represent +/- SE. Kruskal-Wallis *H*-test followed by a Mann-Whitney *U*-test, \*\*\**P*<0.001. Area measurements was performed using segmentation image analysis using ImageJ software. The seed shape was assimilated to its circularity, a circularity of 1 corresponding to a perfect round shape of the seed. (c) Overview of the experimental approach used the segmentation and shape parameters evaluation of the embryos. Row picture (top) and segmented picture (bottom) are illustrated with magnifications of two significant seeds. The same methodology was applied for whole seed segmentation. (d) Embryo area measurements on mature flax embryos. Values are means of four biological replicates, each of thirty seeds. Error bars represent +/- SE. Kruskal-Wallis *H*-test followed by a Mann-Whitney *U*-test, \*\*\**P*<0.001. (e) Seed circularity measurements on mature flax embryos. Values are means of four biological replicates, each of thirty seeds. Error bars represent +/- SE. Kruskal-Wallis *H*-test followed by a Mann-Whitney *U*-test, \*\*\**P*<0.001. The embryo shape was assimilated to its circularity, a circularity of 1 corresponding to a perfect round shape of the embryo. (f-g) Rheological properties analysis of the mature flax seed surface using micro-indentation corresponding to the elastic moduli (f) and hardness (g) measurements. Values are means of thirteen biological replicates with twenty measurements on each (n=260 measurements per RIL). Error bars represent +/- SE. Kruskal-Wallis *H*-test followed by a Mann-Whitney *U*-test, \**P*<0.05, \*\*\**P*<0.001 [20]. (h) Schematic representations of seed and embryo size and shape in the three RILs. These schematic representations don’t account for the actual size and shape of the flaxseed, or even the size of the embryo. The lateral parts of the seeds highlighted in orange and red illustrate the intensity of the mechanical forces exerted by the embryo on the SCE.

It is generally accepted that proliferation of endosperm cells and the regulated control of the seed coat extension play a key role in the mechanisms governing the final size in angiosperms [48,92]. It has also been proposed that endogenous mechanical forces are perceived by a specific cell layer in the seed coat, leading to cell wall thickening which impacts on the seed coat extension and, consequently, on the seed size [50,93,94]. By using micro-indentation microscopy experiments at the seed coat surface - top of the MSCs - differences were revealed in the rheological properties of the RILs seeds (Figure 8f and g). RILs 283 and 44 appeared softer than RIL 80 (Figure 8f) while RIL 283 appeared less plastic than RIL 80 and 44 (Figure 8g).

The Figure 8h shows a diagram illustrating the combination of data from Figures 8a to g. In this diagram, the embryo expansion occurring in RILs 44 and 80 seeds would lead to apply a pressure higher in 80 than in 44, according to the transverse axis. This pressure could locally modify the cell wall properties of, at least, one outer integument specific cell layer. This modification, in turn could have an impact on the physical properties of the seed coat surface resulting in the different release of mucilage observed for the three RILs.

Considering this model and the differences in transcriptional regulations observed between the RILs seeds, it can be hypothesized that pressure exerted by the embryo onto the SCE might regulate, in turn, the expression of some genes involved in the final steps of the mucilage-related processes, in GA- and FA-related metabolisms, and in the directional seed growth.

## CONCLUSION

### A global model for mucilage biosynthesis, oil production and seed growth in flax

The overview of the results, as outlined in Figure 9, shows a seed coat-specific transcriptional regulation of the mucilage chemical composition from early to mid-stage of seed development. These regulations correlate with the use of main carbon sources in the seed coat. In the embryo, however, RIL 44 is close to RIL 80 for C18:2 FA and C18:3 FA from 10 to 25 DPA but gets closer to RIL 283 for C18:2 FA from 25 DPA to mature seed, although no major differential use of photosynthate have been observed (Figure 5f and Figure 7a,b). It has been shown in Arabidopsis that no symplastic connections exist between seed tissues from early to mature embryo, although the mature embryo forms a single symplastic domain [47,88]. Accordingly, our results suggest the existence of independent regulations of both mucilage and FA biosynthesis pathways from early (10 DPA) to mid-stage of seed development. Once the mucilage-related processes established, it is hypothesized that the mechanical pressure exerted by the embryo on the SCE induces seed coat-specific transcriptomes modulations that affect, in turn, the seed oil and FA composition in the embryo, as observed for RIL 44 (Figure 5b-f). Interestingly, in Arabidopsis, extension in time of WRINKLE1 (WRI1) expression in the mid-phase of seed development increases seed oil content [95]. These results also suggest a likely role for GA metabolism in modulating the seed size and shape but also the seed oil and FA composition in response to mechanical stresses (Figure 7, 9, Supplemental Figure S5).

**Figure 9.**
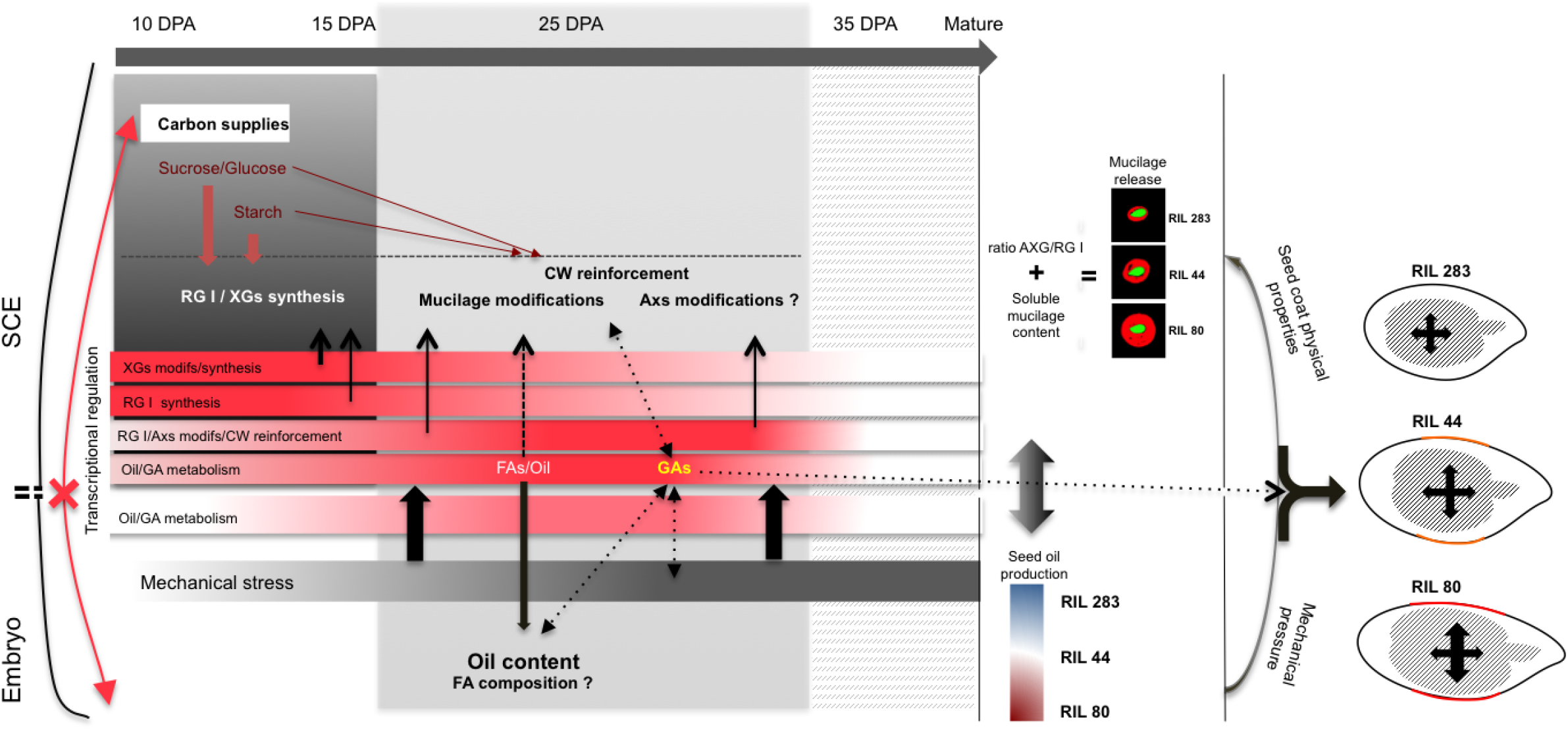
Simplified conceptual model of the possible interactions between the mucilage, FA and GA metabolisms during flax seed tissues development. The most relevant results of this study from our multidisciplinary approach are shown in the diagram, including carbon allocation and mechanical forces from the pressure exerted by the embryo on the SCE. The red arrows indicate that not more carbon allocations between both tissues through the symplasmic pathways are not supposed to exist anymore at 10 DPA. The most relevant tissues- and development stage-specific events are merged in the dark grey and grey boxes. The color code of the red bars indicates the importance of the transcriptional regulation processes for each specific biosynthetic pathway. Phenotypes of the RILs used in this study and corresponding to the mucilage-released as well as FAs contents and the seed size and shape are reported at the end of the seed development (right).

### Limitations of the proposed model

The model proposed here puts emphasis on a positive relationship between the mechano-chemical regulation of the release of mucilage, the soluble mucilage content and the seed oil metabolism. However, the analysis of a number of North American linseed cultivars from the world flax seed collection have not revealed any correlations between the mucilage yield/viscosity (mucilage indicator value; MIV) and the oil content, protein content, seed colour and the thousand seed weight (TSW) [44,45]. The yield of the soluble early-released mucilage was also found as negatively correlated to the vegetative period and the weight of one thousand seeds [96], whereas comparison analysis between brown and yellow flax seeds revealed lower seed weight and higher mucilage production for brown seeds [45]. RILs 283 and 44, which release less soluble mucilage compared with RIL 80, exhibit a lower seed weight (Supplemental Figure S3). This suggests a strong heterogeneity in the joint regulations of the flax mucilage biosynthesis pathways and other seed traits. Integration of our results (Figure 9) suggests that in-depth analyses of the relationships between number of seed traits, including mucilage, seed oil and seed shape, require careful considerations of their complex spatio-temporal transcriptional regulations and insights into the chemical composition of the mucilage [17,74]. In Arabidopsis, a loss of GLABRA2 (GL2) activity in the seed coat leads to reduce mucilage production and enhance seed oil content [33]. It has been hypothesized that the sucrose excess of the mucilage biosynthesis could be transferred in the embryo for contributing to the oil production during the early stages of seed development [33,47]. We could demonstrate the seed coat specific use of carbon supplies and transcriptional regulations from 10 to 25 DPA, however, we cannot exclude such a similar carbon allocation between seed tissues during the very early stages of development, i.e. before 10 DPA, as well as carbon allocation through apoplastic pathways during development from 10 to 25 DPA.

### Strategies to increase seed oil contents

Since these results suggest a plausible correlation between the mucilage content and release and the seed oil production, plant breeding programs focusing on improving seed yield and quality might have to integrate analysis of the seed coat mucilage as a new available breeding way for number of myxospermous species. Classical strategies for the development of transgenic plants with enhanced seed oil levels require genetic engineering of genes encoding enzymes involved in the flow of carbon through modification of one or multiple genes [97,98]. Thus, it is suggested that genetic engineering of a variety of genes involved in the mucilage, seed oil and the GA signaling pathways and especially expressed in the seed coat in the mid-stage of seed development could represent another possible strategy to increase oil content of the flaxseeds.

## SUPPLEMENTAL DATA

**Figure S1:** Phenotyping of the 3 selected RILs with contrasting mucilage release and for the analyze of the shapes of the seed by ImageJ.

a. Mucilage released of the three RILs. Bar = 1,5 cm.

b. Seed size of the three RILs. Bar = 5 mm.

**Figure S2**: Identification of the 3 selected RILs displaying contrasting phenotypes for seed oil content and mucilage release.

Frequency histogram of the recombinant inbred lines (RILs) population screened for seed oil content. Quantification was performed in mature seeds from 186 field-growth RILs (at F10 generation) produced by crossing between Oliver and Viking cultivars. Red dashed lines show seed oil contents found for the RILs selected in this study (283, 44 and 80) for which mucilage phenotypes are shown in the pictures at the top right.

Bar = 3 mm.

**Figure S3**: Seed traits description of the selected RILs. Seed traits description of the selected RILs.

a. Seed coat colour of the three selected RILs. Bar = 8 mm.

b. Relative size of a representative flax seed embryo at 10 DPA. The green embryo can be seen through the transparent seed coat. Bar = 1 mm.

c. Seed weight of the three selected RILs. Error bars represent +/- SE. Kruskal-Wallis *H*-test followed by a Mann-Whitney *U*-test, \*\*\**P*<0.001. Data are means of three biological replicates.

**Figure S4**: HPLC-UV analysis of the lignan content of the SDG-HMG complex.

The quantification of the main lignan content was performed on mature seed from selected RILs. Error bars represent +/- SE (n = 10 to 18 biological replicates). Kruskal-Wallis *H*-test followed by a Mann-Whitney *U*-test, \**P*<0.05.

SDG(+), Secoisolariciresinol diglucoside; FAG, ferulic acid glucoside; CAFG, caffeic acid glucoside; CAG, coumaric acid glucoside.

**Figure S5**: Clustering of the most discriminant genes in pairwise comparisons analysis of RILs in the mid-stage of seed-coat development.

The three clusters correspond to the pairwise comparisons between RILs 44 and 283 (a), RILs 80 and 283 (b) and RILs 80 and 44 (C). Each number associated to the cluster header represents the number of unique gene and overlapping genes for the corresponding pairwise comparison. Genes highlighted in blue are potentially involved in the cell wall and mucilage-related processes. Genes highlighted in red are involved in the seed coat oil and FA metabolisms, while genes highlighted in green correspond to those involved in the GA metabolism. Red dashed lines are positive (>2) and negative (<-2) fold-change thresholds.

**Figure S6**: Analyses of the distribution of the flax genes between the 3 RILs according to their respective functions in the mucilage-related events, across tissues and during seed development. The histogram shows the number of unique genes from pairwise comparison analysis (three-ways ANOVA) of the three RILs. Genes are classified in five different categories according to their roles based on literature.

**Figure S7**: Analysis of the rheological properties of the flax seed surface between the RILs 80 and 283 using micro-indentation experiments.

Boxplots show measurements of the relative elasticity (elastic moduli) at the radial and distal cell wall surfaces of the MSCs for both the RILs 80 and 283. As previously demonstrated for RIL 44 (Miart *et al*., 2019), any statistical differences (Kruskal-Wallis *H*-test followed by a Mann-Whitney *U*-test) were found between measurements on both localisations for the two RILs. Values are means of thirteen biological replicates with twenty measurements on each (n=260 measurements per RIL). Error bars represent +/- SE.

**Table S1**: Discriminating metabolites identified by 1D 1H NMR in the three RILs.

## ACKNOWLEDGMENTS

We thank Helen North (IJPB, INRAE Versailles-Grignon, France) and Georges Haughn (University of British Columbia, Canada) for their critical encouragement, comments and advice. We also thank Jamila Henchi and all the staff of the BIOPI laboratory (University of Picardie Jules Verne) for their help in harvesting flax seeds in the greenhouse and Lorine Tonnelet (University of Cergy-Pontoise, France).

## AUTHORS CONTRIBUTON

F.Me., L.G. and O.V.W. conceived the research project. F.Mi., J-X.F., R.M., G.M., C.W., M.L-P., R.R., L.G., O.V.W., N.D., L.D., A.B., B.T., K.P. and F.Me. designed the experiments and interpreted the results. N.D. and A.B. carried out the micro-indentation experiments and analysed the results. G.M. and C.W. carried out transcriptomic experiments. F.Mi., F.F., H.D., S.B., K.P., dissected the seeds. F.Mi. carried out all experiments and data analysis except for micro-indentation and wrote the paper assisted by L.G. O.V.W., K.P. and F.Me. All authors discussed the results.

