## Supplementary material for "INTEGUMENT-SPECIFIC TRANSCRIPTIONAL REGULATION IN THE MID-STAGE OF FLAX SEED DEVELOPMENT INFLUENCES THE RELEASE OF MUCILAGE AND THE SEED OIL CONTENT": Figure S1, Figure S2, Figure S3, Figure S4, Figure S5, Figure S6, Table S1

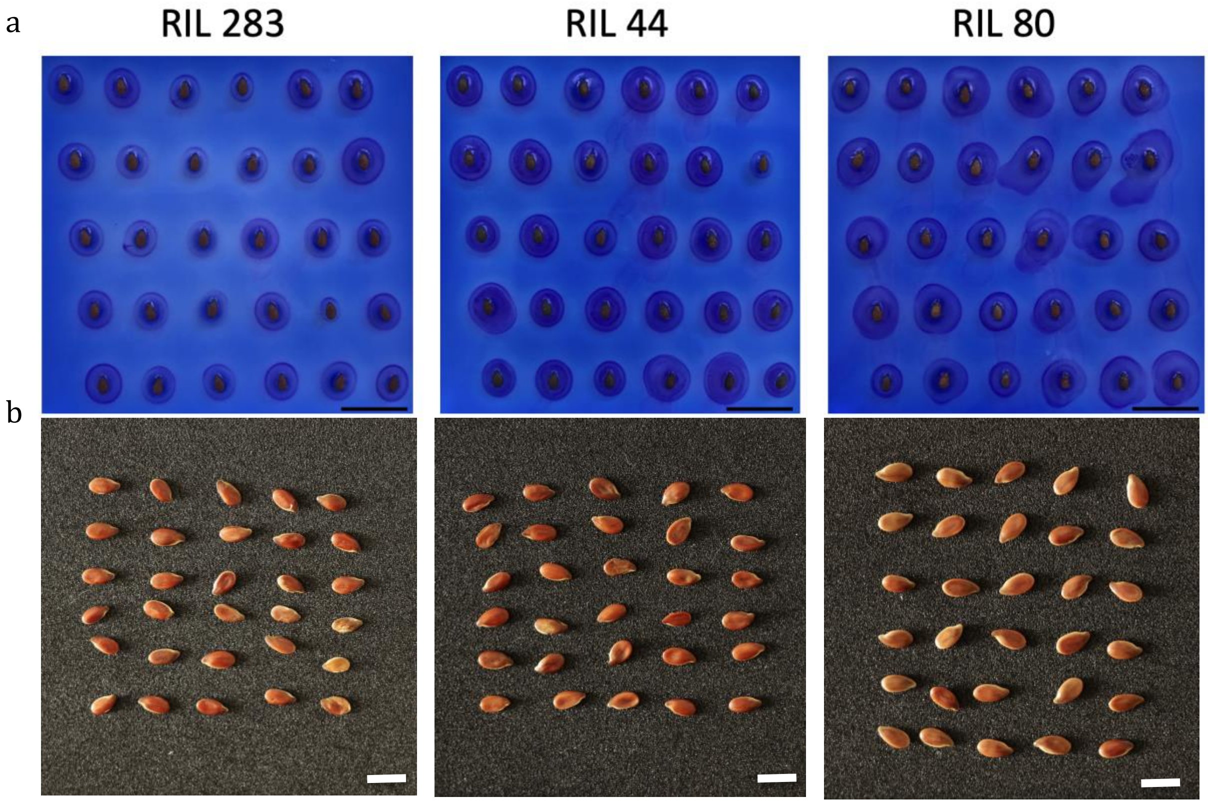


**Figure S1:** Phenotyping of the 3 selected RILs with contrasting mucilage release and for the analyze of the shapes of the seed by ImageJ.

a. Mucilage released of the three RILs. Bar = 1,5cm.

b. Seed size of the three RILs. Bar = 5 mm.

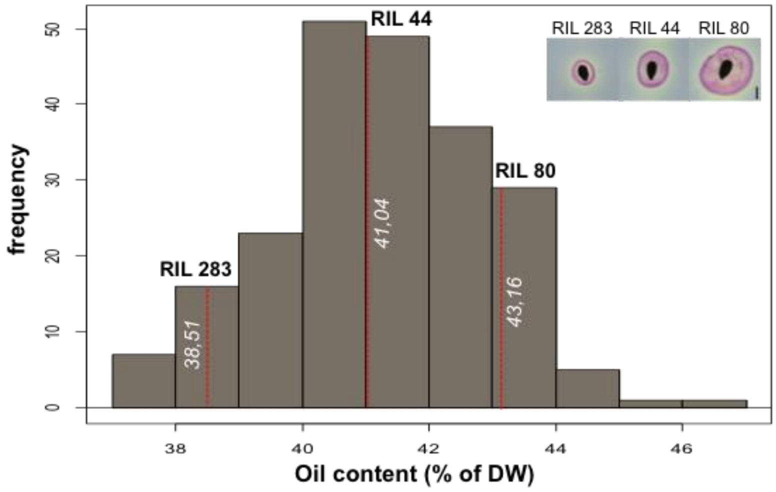


**Figure S2**: Identification of the 3 selected RILs displaying contrasting phenotypes for seed oil content and mucilage release.

Frequency histogram of the recombinant inbred lines (RILs) population screened for seed oil content. Quantification was performed in mature seeds from 186 field-growth RILs (at F10 generation) produced by crossing between Oliver and Viking cultivars. Red dashed lines show seed oil contents found for the RILs selected in this study (283, 44 and 80) for which mucilage phenotypes are shown in the pictures at the top right.

Bar = 3 mm.

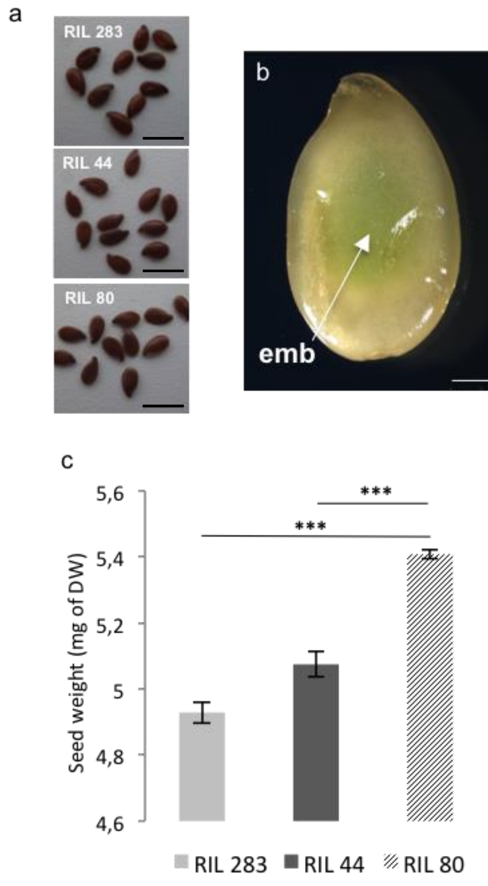


**Figure S3**: Seed traits description of the selected RILs.

a. Seed coat colour of the three selected RILs. Bar = 8 mm.

b. Relative size of a representative flax seed embryo at 10 DPA. The green embryo can be seen through the transparent seed coat. Bar = 1 mm.

c. Seed weight of the three selected RILs. Error bars represent +/- SE. Kruskal-Wallis *H*-test followed by a Mann-Whitney *U*-test, ****P*<0.001. Data are means of three biological replicates.

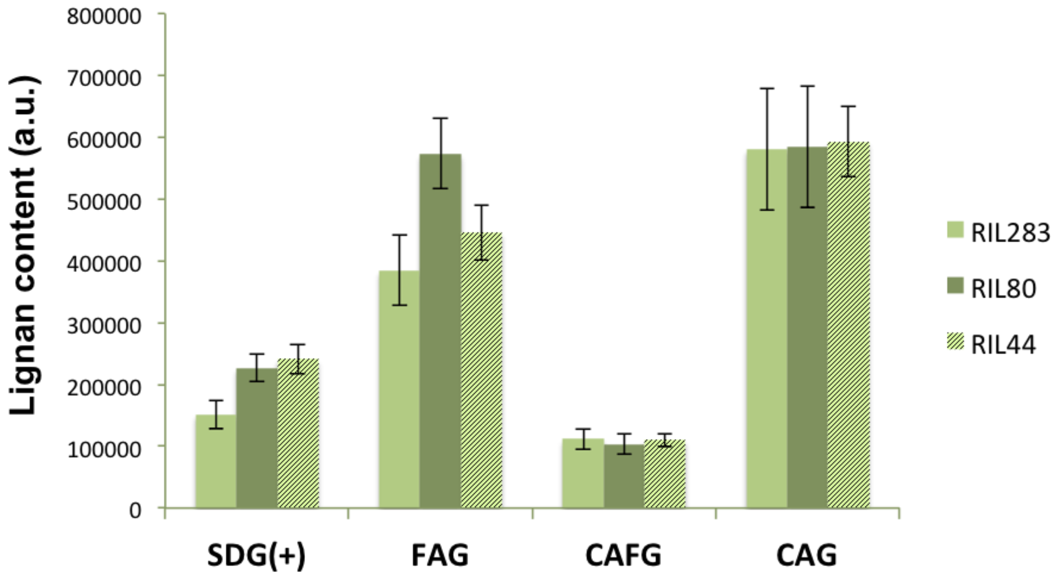


**Figure S4**: HPLC-UV analysis of the lignan content of the SDG-HMG complex.

The quantification of the main lignan content was performed on mature seed from selected RILs. Error bars represent +/- SE (n = 10 to 18 biological replicates). Kruskal-Wallis *H*-test followed by a Mann-Whitney *U*-test, **P*<0.05.

SDG(+), Secoisolariciresinol diglucoside; FAG, ferulic acid glucoside; CAFG, caffeic acid glucoside; CAG, coumaric acid glucoside.

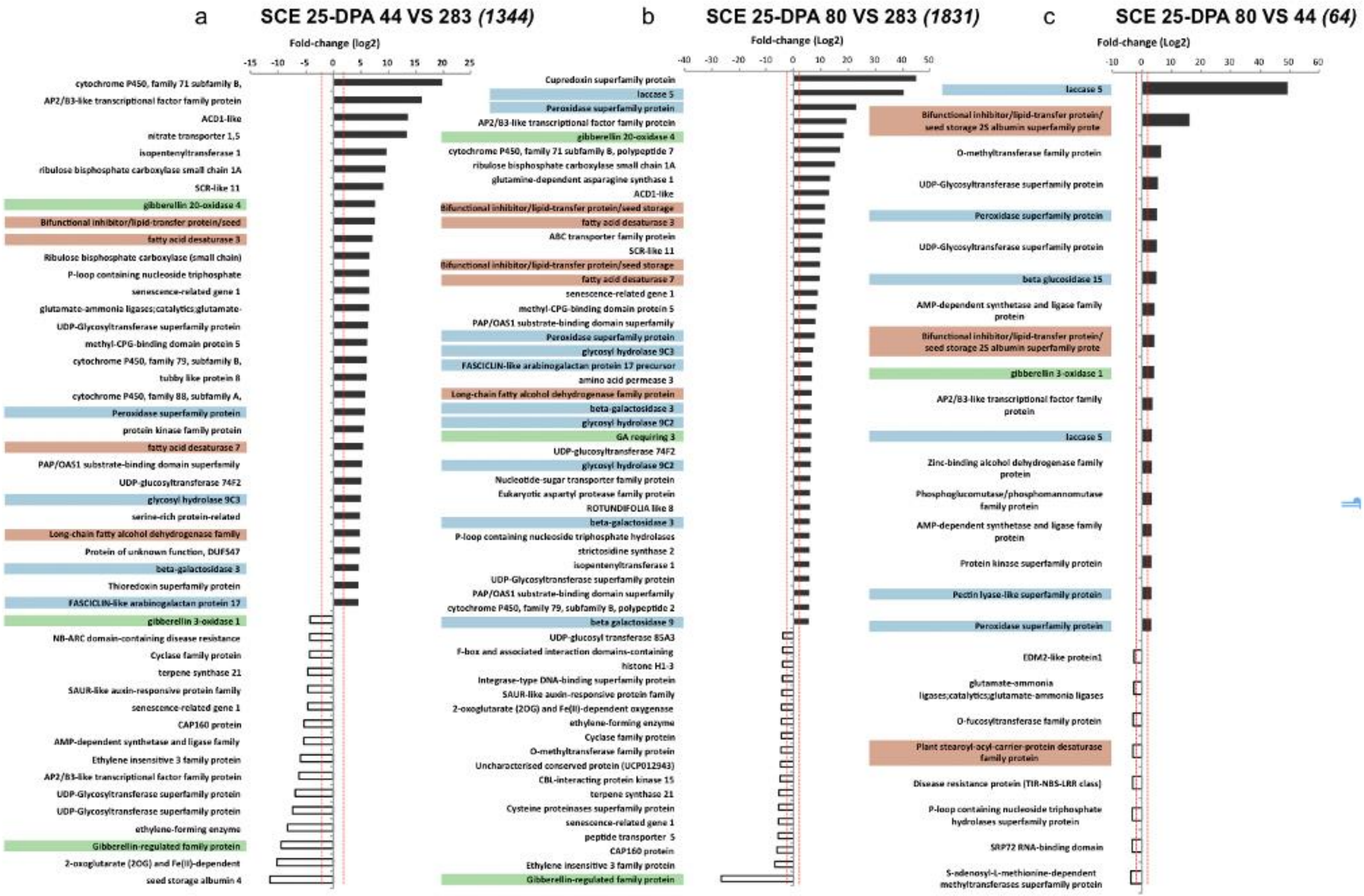
a
SCE 25-DPA 44 VS 283 (1344)
Fold-change (log2)
-15 -10 -5 0 5 10 15 20 25
cytochrome P450, family 71 subfamily B,
AP2/B3-like transcriptional factor family protein
ACD1-like
nitrate transporter 1,5
isopentenyltransferase 1
ribulose bisphosphate carboxylase small chain 1A
SCR-like 11
gibberellin 20-oxidase 4
Bifunctional inhibitor/lipid-transfer protein/seed
fatty acid desaturase 3
Ribulose bisphosphate carboxylase (small chain)
P-loop containing nucleoside triphosphate
senescence-related gene 1
glutamate-ammonia ligases;catalytics;glutamate-
UDP-Glycosyltransferase superfamily protein
methyl-CPG-binding domain protein 5
cytochrome P450, family 79, subfamily B,
tubby like protein 8
cytochrome P450, family 88, subfamily A,
Peroxidase superfamily protein
protein kinase family protein
fatty acid desaturase 7
PAP/OAS1 substrate-binding domain superfamily
UDP-glucosyltransferase 74F2
glycosyl hydrolase 9C3
serine-rich protein-related
Long-chain fatty alcohol dehydrogenase family
Protein of unknown function, DUF547
beta-galactosidase 3
Thioredoxin superfamily protein
FASCICLIN-like arabinogalactan protein 17
gibberellin 3-oxidase 1
NB-ARC domain-containing disease resistance
Cyclase family protein
terpene synthase 21
SAUR-like auxin-responsive protein family
senescence-related gene 1
CAP160 protein
AMP-dependent synthetase and ligase family
Ethylene insensitive 3 family protein
AP2/B3-like transcriptional factor family protein
UDP-Glycosyltransferase superfamily protein
UDP-Glycosyltransferase superfamily protein
ethylene-forming enzyme
Gibberellin-regulated family protein
2-oxoglutarate (2OG) and Fe(II)-dependent
seed storage albumin 4
b
SCE 25-DPA 80 VS 283 (1831)
Fold-change (Log2)
-40 -30 -20 -10 0 10 20 30 40 50
Cupredoxin superfamily protein
laccase 5
Peroxidase superfamily protein
AP2/B3-like transcriptional factor family protein
gibberellin 20-oxidase 4
cytochrome P450, family 71 subfamily B, polypeptide 7
ribulose bisphosphate carboxylase small chain 1A
glutamine-dependent asparagine synthase 1
ACD1-like
Bifunctional inhibitor/lipid-transfer protein/seed storage
fatty acid desaturase 3
ABC transporter family protein
SCR-like 11
Bifunctional inhibitor/lipid-transfer protein/seed storage
fatty acid desaturase 7
senescence-related gene 1
methyl-CPG-binding domain protein 5
PAP/OAS1 substrate-binding domain superfamily
Peroxidase superfamily protein
glycosyl hydrolase 9C3
FASCICLIN-like arabinogalactan protein 17 precursor
amino acid permease 3
Long-chain fatty alcohol dehydrogenase family protein
beta-galactosidase 3
glycosyl hydrolase 9C2
GA requiring 3
UDP-glucosyltransferase 74F2
glycosyl hydrolase 9C2
Nucleotide-sugar transporter family protein
Eukaryotic aspartyl protease family protein
ROTUNDIFOLIA like 8
beta-galactosidase 3
P-loop containing nucleoside triphosphate hydrolases
strictosidine synthase 2
isopentenyltransferase 1
UDP-Glycosyltransferase superfamily protein
PAP/OAS1 substrate-binding domain superfamily
cytochrome P450, family 79, subfamily B, polypeptide 2
beta galactosidase 9
UDP-glucosyl transferase 85A3
F-box and associated interaction domains-containing
histone H1-3
Integrase-type DNA-binding superfamily protein
SAUR-like auxin-responsive protein family
2-oxoglutarate (2OG) and Fe(II)-dependent oxygenase
ethylene-forming enzyme
Cyclase family protein
O-methyltransferase family protein
Uncharacterised conserved protein (UCP012943)
CBL-interacting protein kinase 15
terpene synthase 21
Cysteine proteinases superfamily protein
senescence-related gene 1
peptide transporter 5
CAP160 protein
Ethylene insensitive 3 family protein
Gibberellin-regulated family protein
c
SCE 25-DPA 80 VS 44 (64)
Fold-change (Log2)
-10 0 10 20 30 40 50 60
laccase 5
Bifunctional inhibitor/lipid-transfer protein/ seed storage 2S albumin superfamily prote
O-methyltransferase family protein
UDP-Glycosyltransferase superfamily protein
Peroxidase superfamily protein
UDP-Glycosyltransferase superfamily protein
beta glucosidase 15
AMP-dependent synthetase and ligase family protein
Bifunctional inhibitor/lipid-transfer protein/ seed storage 2S albumin superfamily prote
gibberellin 3-oxidase 1
AP2/B3-like transcriptional factor family protein
laccase 5
Zinc-binding alcohol dehydrogenase family protein
Phosphoglucomutase/phosphomannomutase family protein
AMP-dependent synthetase and ligase family protein
Protein kinase superfamily protein
Pectin lyase-like superfamily protein
Peroxidase superfamily protein
EDM2-like protein1
glutamate-ammonia ligases;catalytics;glutamate-ammonia ligases
O-fucosyltransferase family protein
Plant stearoyl-acyl-carrier-protein desaturase family protein
Disease resistance protein (TIR-NBS-LRR class)
P-loop containing nucleoside triphosphate hydrolases superfamily protein
SRP72 RNA-binding domain
S-adenosyl-L-methionine-dependent methyltransferases superfamily protein

**Figure S5**: Clustering of the most discriminant genes in pairwise comparisons analysis of RILs in the mid-stage of seed-coat development.

The three clusters correspond to the pairwise comparisons between RILs 44 and 283 (a), RILs 80 and 283 (b) and RILs 80 and 44 (C). Each number associated to the cluster header represents the number of unique gene and overlapping genes for the corresponding pairwise comparison. Genes highlighted in blue are potentially involved in the cell wall and mucilage-related processes. Genes highlighted in red are involved in the seed coat oil and FA metabolisms, while genes highlighted in green correspond to those involved in the GA metabolism. Red dashed lines are positive (>2) and negative (<-2) fold-change thresholds.

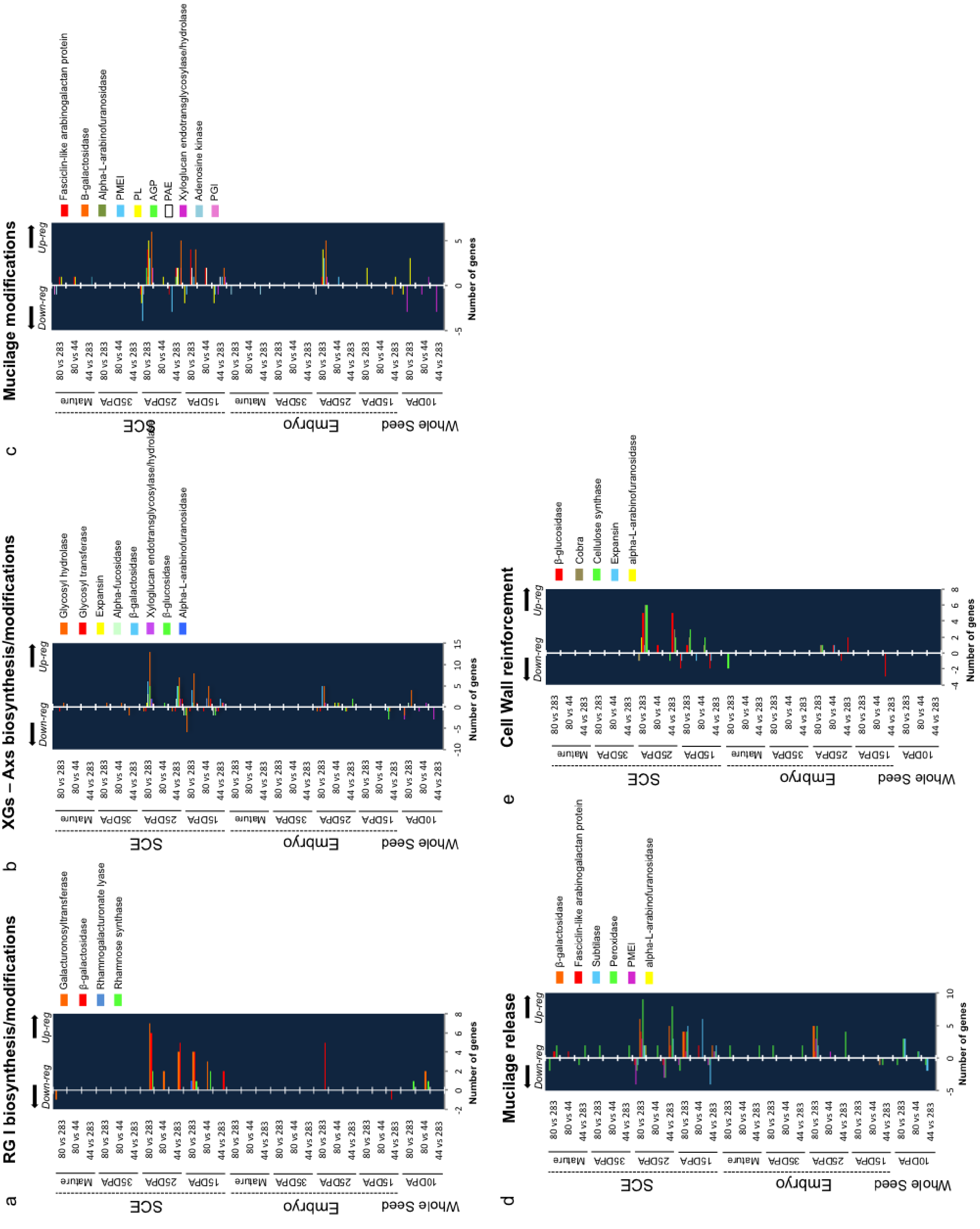


**Figure S6**: Analyses of the distribution of the flax genes between the 3 RILs according to their respective functions in the mucilage-related events, across tissues and during seed development.

The histogram shows the number of unique genes from pairwise comparison analysis (three-ways ANOVA) of the three RILs. Genes are classified in five different categories according to their roles based on literature.

**Table S1**: Discriminating metabolites identified by 1D 1H NMR in the three RILs.

| *Compounds* | *Integrated area* | | *Chemical shift δH and multiplicity on 1H spectrum (coupling constant J) of the peak used for calculation of the areas* |
|---|---|---|---|
| L-Valine | 1.012 | 1.0 | CH3, 1,011 ppm, d (7 Hz) |
| L-Alanine | 1.502 | 1.475 | CH3, 1,49 ppm, d (7.23 Hz) |
| Linamarin | 4.572 | 4.56 | CH, 4.57 ppm, d (7.81 Hz) |
| Neolinustatine | 4.483 | 4.47 | CH, 4.48 ppm, d (7.84 Hz) |
| Linustatine | 4.45 | 4.442 | CH, 4.45 ppm, d (7.71 Hz) |
| L-Asparagine | 2.95 | 2.932 | CH2, 2.95 ppm, dd (16.92; 4,03 Hz) |
| Ethanolamine | 3.128 | 3.1 | CH2, 3.12 ppm, m |
| Choline | 3.221 | 3.2125 | CH3, 3.22 ppm, s |
| Raffinose D(+) | 4.96 | 4.92 | CH, 4.94 ppm, d (3.71 Hz) |
| Sucrose | 4.165 | 4.148 | CH, 4.16 ppm, d (8.61 Hz) |
| L-Tryptophane | 7.29 | 7.274 | CH, 7.28 ppm, s |
| Trigonelline | 9.17 | 9.14 | CH, 9.155 ppm, s |
| Formic acid | 8.485 | 8.46 | CH, 8.47 ppm, s |
| L(+)-Tartaric-acid | 4.335 | 4.312 | CH, 4.32 ppm, s |
| L-Aspartic-acid | 2.679 | 2.652 | CH2, 2.67 ppm, dd (17.3; 8.90 Hz) |
| L-Glutamic-acid | 2.436 | 2.397 | CH2, 2.41 ppm, m |
| L-Glutamine | 2.482 | 2.437 | CH2, 2.46 ppm, m |
| D(+)-Glucose | 3.185 | 3.175 | CH, 3.20 ppm, dd (9.30; 7.93 Hz) |
| Glycerol | 3.518 | 3.511 | CH2, 3.515, m |
| Acetate | 1.96 | 1.949 | CH3, 1,955 ppm, s |
